# Structure and sequence evolution in the pennycress (*Thlaspi arvense*) pangenome

**DOI:** 10.1101/2025.09.26.678720

**Authors:** Kevin A. Bird, Joanna L. Rifkin, Chloee M. McLaughlin, Avril M. Harder, Pawan Basnet, Ella Katz, Tomáš Brůna, Kerrie Barry, LoriBeth Boston, Christopher Daum, Jie Guo, Anna Lipzen, Christopher Plott, Jerry W. Jenkins, Rachel Walstead, Shanmugam Rajasekar, Jayson Talag, Katherine Frels, Kathleen Greenham, Shelby Ellison, Jane Grimwood, Jeremy Schmutz, Patrick P. Edger, J. Chris Pires, John T. Lovell, Daniel Kliebenstein

## Abstract

- Eukaryotic genomes harbor many forms of variation, including nucleotide diversity and structural polymorphisms, which experience natural selection and contribute to genome evolution and biodiversity. However, harnessing this variation for agriculture hinges on our ability to detect, quantify, catalog, and utilize genetic diversity.
- Here, we explore seven complete genomes of the emerging biofuel crop pennycress (*Thlaspi arvense*) drawn from across the species’s current genetic diversity to catalogue variation in genome structure and content.
- Across this new pangenome resource, we find contrasting evolutionary modes in different genomic regions. Gene-poor, repeat-rich pericentromeric regions experience frequent rearrangements, including repeated centromere repositioning. In contrast, conserved gene-dense chromosome arms maintain large-scale synteny across accessions, even in fast-evolving immune genes where microsynteny breaks down across species but the macrosynteny of gene cluster positioning is maintained.
- Our findings highlight that multiple elements of the genome experience dynamic evolution that conserves functional content on the chromosome scale but allows rearrangement and presence-absence variation on a local scale. This diversity is invisible to classical reference-based approaches and highlights the strength and utility of pangenomic resources. These results provide a valuable case study of rapid genomic structural evolution within a species and powerful resources for crop development in an emerging biofuel crop.

## Introduction

Genetic diversity in crops allows for resilience and sustainability in agricultural systems. In established crops, the DNA sequence variation underlying quantitative genetic variation is leveraged to develop trait combinations and cultivars that are resilient to changing climates, respond to novel pests and pathogens, and minimize the environmental impact of agriculture. There has also been considerable effort to develop new crops from wild species that already harbor stress tolerance and novel end use trait variation (e.g. seed oil chemistry). However, such *de novo* domestication is typically constrained by limited genetic variation within traits needed for efficient growth and sufficient yield in agricultural systems (e.g. loss of shattering, low seed dormancy, etc.). These constraints can be circumvented by genome editing or mutant screens (Lemmon et al., 2018; Zsögön et al., 2018; Gasparini et al., 2021), but these approaches require species amenable to transformation with genomes that are tractable for investigations of genetic variation underlying the traits of interest (e.g. loss-of-function, gain-of-function, dosage change, etc). These requirements become more onerous with greater evolutionary distance to model systems (Roeder et al., 2025). Thus, improvement of domesticated species, and successful development of new crop species, hinges on our ability to detect, quantify, catalog, and deploy genetic diversity.

Genome sequences and associated resources facilitate surveys of genetic diversity, identification of valuable germplasm, and discovery of causal loci that underlie traits of interest in crops and crop wild relatives, with the ultimate goal of utilizing well characterized genetic variation to develop new elite cultivars. However, the success of these transfers depends on the genomic context of sequence variation that underlies traits of interest. In an ideal scenario, traits are unlinked and in high-recombination regions allowing desirable alleles to be stacked without interference from deleterious variants. However, gene density and recombination rate vary predictably in most plant genomes, sorting most sequence into two ‘compartments’: ‘pericentromeres’ are low-recombination highly repetitive regions adjacent to centromeres, while ‘chromosome arms’ typically harbor most of the genes and recombination events. Such covariance between chromosome architecture and recombination landscape impacts the evolutionary dynamics and accessibility of genetic diversity to selection agents. For example, pericentromeric sequences often have weaker signatures of purifying selection than chromosome arms, likely due to lower recombination rates and increased effects of linkage drag (Chen et al., 2022). Such relaxed selection pressure means that pericentromeres often harbor greater sequence and structural diversity than subtelomeric chromosome arms (Kawabe et al., 2004; Hall et al., 2006; Flowers et al., 2012), sometimes to such extent that they are the proximate cause of reproductive barriers between species (Ma and Zhu, 2025). Despite millennia of selection and intercrossing, most domesticated species exhibit variable degrees of reproductive isolation, which can constrain efforts to introgress traits between highly diverged gene pools (e.g. sorghum, rice, etc.). Therefore, it is likely that contemporary efforts by breeders to domesticate plant species and develop new crops will confront challenges related to reproductive isolation, possibly due to structural evolution among pericentromeres.

Pennycress (*Thlaspi arvense* L.; 2n=2x=14), an annual weedy species in the *Brassicaceae* family native to Eurasia, offers an excellent system to explore functional and evolutionary consequences of pangenomic variation during *de novo* domestication. Several attractive agronomic and genomic attributes have made pennycress a target for domestication and improvement for use as a winter cover crop, oil feedstock, and biofuel source (Boateng et al., 2010; Moser, 2012; Fan et al., 2013). Agronomically, the winter hardiness and short life cycle of pennycress make it well-suited to double-crop systems that can increase productivity and foster sustainable agricultural practices by reviving soil health, reducing erosion, providing pollinator resources, and improving land use efficiency (Phippen and Phippen, 2012; Del Gato et al., 2015; Johnson et al., 2015; Weyer et al., 2019; 2021; Basnet and Ellison, 2023). Pennycress has a medium-sized diploid genome (∼526Mb based on flow cytometry; Lysak et al., 2009) and a predominantly one-to-one gene ratio with the model plant *Arabidopsis thaliana* (Arabidopsis), both of which aid domestication and improvement efforts by accelerating identification of candidate orthologs with functional effects on key traits (Chopra et al., 2018; 2020; Roeder et al., 2025). Both classical breeding and targeted gene editing efforts are further bolstered by the predominantly self-fertilizing habit of pennycress and its amenability to floral dip transformation (Mulligan and Kevan, 1973; McGinn et al., 2019). Despite these advantages enabling *de novo* domestication, pennycress’s limited genetic diversity and susceptibility to biotic pests and pathogens such as soybean cyst nematode, aphids, black rot, black spot disease, and clubroot disease are persistent barriers to widespread adoption (Basnet and Ellison, 2023).

Research and development of pennycress has greatly benefited from the recently improved reference genome (Dorn et al., 2015; Garcia Navarette et al., 2022; Nunn et al., 2022), which has supported short-read resequencing of pennycress diversity panels to characterize genetic diversity in the species across its range. Populations from Western Europe and North America have low levels of genetic distance, as expected given that pennycress is believed to have been introduced to North America by European immigration within the last 250 years (Frels et al., 2019; Nunn et al., 2022). In its native range, pennycress exhibits minimal population structure between Western European and East Asian populations. However, multiple studies have identified populations in Armenia as genetically distinct from all other pennycress populations, showing high levels of genetic distance and unidirectional gene flow out of but not into Armenia (Frels et al., 2019; Nunn et al., 2022; Contreras-Garrido et al., 2024; Wu et al., 2025). The single reference genome also enabled transposable element surveys, creating a list of transposon insertion polymorphisms that may be harnessable by scientists and breeders to accelerate crop improvement (Nunn et al., 2022; Contreras-Garrido et al., 2024). While existing resources have identified candidate resistance loci through genome-wide association studies (Galanti et al., 2024), characterizing resistant accessions and incorporating them into breeding programs requires a deeper understanding of genetic diversity in the species.

Improvements in sequencing technology have reduced the cost and effort needed to construct reference-quality genomes, particularly in plants, which have historically been difficult to sequence because of whole-genome duplications, extensive repeats, and chromosomal rearrangements. Thanks to these improvements, collections of diverse reference genomes for a single species, known as pangenomes, are now available more broadly across the tree of life. Pangenome datasets enable better surveys of genetic variation, revealing previously hidden gene presence-absence variation and the structural variation that shapes adaptation and gene flow across populations (Bayer et al., 2020; Katz et al., 2021; L. Zhang et al., 2021, Zhou et al., 2022; Schreiber et al., 2024). These pangenomic catalogs of sequence variation improve our ability to identify the genetic bases of traits and apply molecular breeding approaches, like *de novo* domestication (Gasparini et al., 2021; Zhou et al., 2022; S. Jin et al., 2023; Z. Liu et al., 2024). This is particularly the case for complex loci such as the NOD-Like Receptor (NLR) pathogen resistance genes, which often undergo rapid duplication and loss (Teasdale et al., 2025).

To support the development of pennycress pangenomic resources, we present seven complete, chromosome-scale genomes sampled across the breadth of genetic diversity of the species. Our pangenomic analyses found a ‘two-speed’ genomic compartmentalization for structural variation. Gene rich chromosome arms exhibited high levels of synteny and low levels of gene Presence/absence variation (PAV) among accessions. In contrast, gene-poor highly repetitive peri- and centromeric regions contained dramatic genomic rearrangement, especially between Armenian and non-Armenian accessions. These genomic rearrangements included centromeric movement within the species, and population genomic analyses suggested these rearrangements may partially limit gene flow among populations. Finally, to support efforts to breed for disease resistance, we developed a catalog of high-confidence NLR genes and identified a subset of NLR loci as highly variable. Comparative genomic analyses revealed that NLR gene clusters tend to be positionally conserved across significant evolutionary distances between species, but rapidly diversify and turn over such that particular NLR genes are unlikely to be orthologous between species.

## Methods

### Plant material, DNA, and RNA extraction

High molecular weight DNA was extracted using the protocol of Doyle and Doyle (1987) with minor modifications. Two grams flash-frozen biomass was ground to a fine powder in a frozen mortar with liquid nitrogen followed by very gentle extraction in 2% CTAB buffer (that included proteinase K, PVP-40 and beta-mercaptoethanol) for 30min to 1h at 50 °C. After centrifugation, the supernatant was transferred to a new tube, treated with 200ul 50mM PSMF for 10 minutes at room temperature then gently extracted twice with 24:1 Chloroform : Isoamyl alcohol. The upper phase was transferred to a new tube and 1/10th volume 3 M Sodium acetate was added, gently mixed, and DNA precipitated with iso-propanol. DNA precipitate was collected by centrifugation, washed with 70% ethanol, air dried for 5-10 minutes and dissolved thoroughly in elution buffer at room temperature followed by RNAse treatment. DNA purity was measured with Nanodrop, DNA concentration measured with Qubit HS kit (Invitrogen), and DNA size was validated by Femto Pulse System (Agilent).

### DNA sequencing and genome assembly

We employed a whole-genome shotgun sequencing strategy for the seven *T. arvense* genomes. Sequencing libraries were constructed for all accessions for PacBio HiFi sequencing using Circular Consensus Sequencing (CCS) mode. We used a Diagenode Megaruptor 3 for DNA shearing, prepared libraries using SMRTbell Template Prep Kit 2.0 kits, and sized our libraries using a SAGE ELF instrument with 1-18kb cassette. Illumina polishing libraries were prepared for all accessions using Illumina TruSeq PCRfree library prep kits (Catalog #20015962). In addition, Omni-C libraries were prepared for extra scaffolding using Dovetail Omni-C kits (Catalog #21005) for MN106 and Ames 32873.

We sequenced our PacBio libraries on the SEQUEL II platform using 1-3 single molecule, real-time (SMRT) cells with V2 chemistry, and our Illumina libraries on the NovoSeq 6000 platform. All sequencing was performed at the HudsonAlpha Institute in Huntsville, AL, US. All accessions were sequenced with PacBio coverage depths between 70.23-88.35x and read lengths averaging between 15,194-18,898bp (Supplementary Data 1). For polishing, all accessions were sequenced with Illumina (2x150, 400bp insert) coverage depths between 49.1x and 54.7x. We also sequenced 2 x 150bp Omni-C libraries for MN106 and Ames 32873.

Genomes were assembled using HiFiAsm v0.16.1-r375 (Cheng et al., 2021; 2022) with Hi-C integration (-h1 and -h2) used for MN106 and Ames 32873. For MN106 and Ames 32873, contigs were aligned and oriented using Hi-C reads with the JUICER pipeline. For all other genomes, contig positions were finalized using a total of 11,295 unique, non-repetitive, non-overlapping 1KB syntenic markers from the version 3.0 MN106 genome aligned to the RACON-polished assembly using BLAT (Kent, 2002). The initial assemblies consisted of between 11 and 23 contigs, which were combined using between 5 and 8 joins into 7 scaffolds. Every genome contained at least one single-contig chromosome, and no chromosome consisted of more than six contigs. No contig breaks were necessary for any of the genomes assemblies. In a final round of polishing, Illumina sequence was used to correct homozygous SNPs and indels with a pipeline composed of bwa mem (v2.2.1; Li and Durbin 2009) and GATK’s UnifiedGenotyper tool (v3.7; McKenna et al., 2010). Fewer than 21 SNPs and 600 indel errors were corrected for all genomes except Tibet 33, in which over 1200 SNPs and over 4300 indels were corrected.

We assessed the completeness of the euchromatic portion of the genomes by aligning *T. arvense* V1 annotated genes to each assembly with BLAT, retaining the longest alternative splicing variant and excluding alignments with <90% identity or <85% coverage. Completeness estimated this way was 97.86 for the highly divergent Ames 32879 genome but between 99.04-99.33 for all other accessions.

### RNA sequencing and genome annotation

For all seven accessions, sequencing libraries were prepared for PacBio Iso-Seq RNA sequencing. We synthesized full-length cDNA with NEBNext Single Cell/Low Input cDNA Synthesis & Amplification Module kit (Catalog #E6421) by amplifying first-strand cDNA with NEBNext High-Fidelity 2× PCR Master Mix (Catalog #M0541) with barcoded cDNA PCR primers for 11-14 cycles. We then purified cDNA using 1× AMPure PB beads (PacBio Catalog #100-265-900) for nonsize selection or BluePippin (Sage Science, Beverly, MA, USA) for 2–10 kb size selection. Sizes were pooled in equimolar ratios according to the PacBio Multiplexing Calculator worksheet (PacBio, Menlo Park, CA, USA). We next end-repaired, A-tailed, and ligated overhang nonbarcoded adaptors to our samples using SMRTbell Express 2.0 kit (PacBio Catalog #100-938-900). PacBio Sequencing primer was annealed to the SMRTbell template library. Sequencing polymerase was bound to samples using Sequel II Binding kit 2.0 (PacBio Catalog #101-789-500). Finally, we sequenced the libraries on a PacBio Sequel IIe [or Revio??] sequencer. Sequencing used SMRT Link 10.2, sample-dependent sequencing primer, 8M v1 SMRT cells, and Version 2.0 sequencing chemistry with 1 × 1,800 sequencing movie run times (PacBio).

We also prepared Illumina sequencing libraries. Plate-based sample prep was performed using Illumina TruSeq-stranded mRNA HT sample prep kits (Catalog #20020595) on PerkinElmer Sciclone NGS robotic liquid handling system (Revvity, Waltham, MA, USA) according to Illumina’s user guide (https://support.illumina.com/sequencing/sequencing_kits/truseq-stranded-mrna.html). We started with 1000 ng per sample and amplified for 8 samples of PCR. Libraries were quantified with KAPA Biosystems next-generation sequencing library qPCR kits (Roche Catalog #07980140001) and run on a Roche LightCycler 480 real-time PCR instrument (Roche Diagnostics, Indianapolis, IN, USA). Samples were sequenced on an Illumina NovaSeq sequencer using NovaSeq XP V1.5 reagent kits (Illumina, San Diego, CA, USA), S4 flowcell, following a 2 × 151 indexed run recipe.

We annotated each genome in two rounds.

#### Round one

In the first round, we generated transcriptome assemblies for each genome using 2 x 150bp Illumina RNA-seq reads with PERTRAN (Lovell et al., 2018), which uses GSNAP (T. D. Wu and Nacu 2010) for genome-guided transcriptome assembly before validating, realigning, and correcting alignments and building a splice alignment graph. We then used a genome-guided correction pipeline to collapse and correct PacBio Iso-Seq CCSs to obtain putative full-length transcripts. We aligned CCS reads to the genome with GMAP (T. D. Wu and Katanabe, 2005; T. D. Wu et al., 2016), corrected small indels in splice junctions, and clustered alignments with all introns >= 95% overlapping. PASA (Haas et al., 2003) was then applied to merge the Illumina-based and PacBio-based transcriptome assemblies.

We then created a *de novo* repeat library from repeats predicted by RepeatModeler2 (Flynn et al., 2020) from the *T. arvense* MN106 v3.0 genome assembly. Repeats were restricted to those with significant protein-coding domains as identified by InterProScan (Jones et al., 2014) with the Pfam (Mistry et al., 2020) and PANTHER (Mi et al., 2019) databases. This library was used to soft-mask each genome with RepeatMasker (Smit et al., 2013-2015).

To identify putative gene loci, we integrated our transcript assembly alignments with EXONERATE (Slater and Birney 2005) alignments of proteins from diverse plants (*Eutrema salsugineum*, *Capsella rubella*, *Arabidopsis thaliana*, *Rorippa islandica*, *Myagrum perfoliatum*, *Caulanthus amplexicaulis, Cleome violacea, Glycine max, Vitis vinifera, Liriodendron tulipifera, Brassica rapa*, *Malcolmia maritima*, *Cleomella arborea*, *Capparis spinosa, Schrenkiella parvula*, *Gossypium raimondii*, *Populus trichocarpa*, *Sorghum bicolor*, *Oryza sativa*, *Beta vulgaris*) and the Swiss-Prot release 2022_04 of eukaryote proteomes to the repeat-soft-masked genome. We allowed up to 2kbp extension on both ends, except when that extended into another locus on the same strand. We used the homology-based predictors FGENESH+ (Salamov and Solovyev 2000), and FGENESH_EST (which uses expressed sequence tags rather than open reading frames to compute splice sites and introns), EXONERATE, PASA assembly ORFs (in-house homology constrained ORF finder), and AUGUSTUS (Stanke et al., 2006) trained on the high-confidence PASA assembly ORFs and with intron hints from short read alignments. We selected the best-scored predictions from each locus using multiple positive factors, which included expressed sequence tags and protein support, and the negative factor of overlap with repeats. We improved these predictions by using PASA to add UTRs, correct splicing, and add alternative transcripts.

We then performed protein homology analysis, comparing the PASA-improved gene models to the diverse proteomes above, resulting in a Cscore (protein BLASTP score ratio to the mutual best hit BLASTP score) and protein coverage (percentage of protein aligned to the best of homologs). Transcripts with Cscore and protein coverage >= 0.5 were retained, as were transcripts covered by ESTs. However, gene models whose CDS overlapped repeats by more than 20% were retained only with Cscore >=0.9 and protein coverage >=0.7. Retained gene models were subjected to Pfam analysis, and models without strong transcriptome and homology support with more than 30% overlap of Pfam TE domains were also removed.

#### Round two

After first-round genome annotation, each genome was hard-masked with its own high-confidence (complete, transcriptome- and homology-supported) gene models. The hard-masked genomes were then aligned using BLASTX and EXONERATE against high-confidence peptides from the other six genomes to make EXONERATE gene predictions. The resulting gene models were scored with BLASTP using homology proteomes. If the new models either a) exhibited higher homology support and were not contradicted by transcriptome evidence or b) were situated at loci without first-round gene models, they were used to replace the original models. We manually removed any gene models that met any of the following criteria: was incomplete, had low homology support and lacked full transcriptome support, had short single exons (<300bp CDS) without protein domains or expression support, or was repetitive and lacked strong homology support.

### Synteny analysis, structural variant identification and pangenome graph construction

We identified general patterns of synteny and gene conservation within our *T. arvense* accessions and in related species in the Brassicaceae using GENESPACE version 1.3.1 (Lovell et al., 2022). For GENESPACE analyses, we identified regions of gene synteny between all seven *T. arvense* accessions and the Phytozome (Goodstein et al., 2012) repository versions of *Arabidopsis thaliana* (v. TAIR10/TAIR11), *A. lyrata* (v. 2.1), and *Brassica rapa* (v. ssp. *trilocularis* R500 v2.1) genomes. Syntenic blocks from GENESPACE and phylogenetic hierarchical orthogroups from Orthofinder v 2.5.5 (Emms and Kelley, 2019) run within GENESPACE were used in downstream analyses.

To identify and classify structural variation, each genome was aligned to the MN106 reference using minimap2 (v2.26; Li 2021) with asm5 preset options. The resulting alignments were filtered in R (v4.4.1; R Core Team) to retain segments ≥1 kb in length with ≥80% sequence identity and MAPQ ≥2, and SyRI (v1.6.3; Goel et al., 2019) was used to classify these alignments as syntenic regions or structural rearrangements. We constructed chromosome-level pangenome graphs with Minigraph-Cactus (v2.9.0; Hickey et al., 2023) using default settings and MN106 as the primary reference. In the last step of graph construction, large unaligned portions (e.g., repeat arrays) of non-primary references are clipped out to produce the final graph. Clipped chromosome graphs were subsequently merged with vg toolkit (v1.59.0; Garrison et al., 2018) and the degree of sequence sharing across haplotypes in the graph was assessed with Panacus (v0.2.3; Parmigiani et al., 2024a). We also calculated 31-mer presence-absence in the genome assemblies to calculate whole-genome sequence growth curves with Pangrowth (Parmigiani et al., 2024b). The same process was repeated for genic sequences and NLRs (see “NLR identification” below).

### Gene ontology (GO) enrichment analysis

We performed Gene Ontology (GO) enrichment analysis using the R package topGO v 2.60.1 (Alexa and Rahnenfuhrer, 2025) and the TAIR Arabidopsis gene library (Reiser et al., 2024). We selected orthogroups from our analysis that included Arabidopsis genes and were either dispensable or unique in the *T. arvense* genomes. We determined enrichment of different biological processes using Fisher’s exact test in topGo.

### Variant calling

Variants for the sequenced reference genomes were called from the Illumina polishing libraries (see DNA sequencing and genome assembly, above). Reads were aligned to the MN106 reference using bwa-mem2 (v2.2.1; Vasimuddin et al., 2019; alignment scripts) and alignments were postprocessed using PicardTools (citation; scripts). Variants were called using bcftools (v1.20; Danecek et al., 2021; scripts) and filtered for minimum and maximum sequencing depth and missing data (scripts).

Genomic DNA was extracted from 95 *T. arvense* samples representing a broad geographic distribution (Supplemental Table 2) using a modified CTAB extraction. DNA was quantified with a Quant-iT™ PicoGreen® dsDNA kit (Life Technologies, Grand Island, NY) and Genotyping-by-Sequencing (GBS) was performed as in (Elshire et al., 2011), at the University of Wisconsin-Madison Biotechnology Center with half-sized reactions. Samples were digested using the *ApeK*I restriction enzyme, barcoded, and pooled for sequencing. Sample libraries were sequenced on an Illumina NovaSeq X Plus sequencer using 2x150bp reads. Samples were demultiplexed with cutadapt (v4.9; Martin 2011) and raw reads were aligned to the MN106 reference genome using bwa-mem2 (v2.21; Vasimuddin et al., 2019). GBS variants were called using bcftools (v1.20; Danecek et al., 2021) and filtered with vcftools (v0.1.16; Danecek et al., 2011) to retain sites with ≤ 30% missingness, a quality score ≥ 30, minimum depth ≥ 5, and maximum depth ≤ 50, resulting in 637,506 sites.

### Principal component analysis

We selected a subset of 163,376 variants genotyped in both the GBS and the reference polishing libraries to perform principal component analysis using the R package SNPRelate (Zheng et al., 2012, supplementary material: PennycressPCA.R). We further filtered these variants for minor allele frequency and removed closely linked variants (LD = 0.95), leaving 18,375 variants.

### Fst analysis

We used a machine-learning approach to classify the genome into ‘chromosome arms’, ‘pericentromere’, and ‘centromere’ based on their genic and repetitive element content. The boundaries of these regions were then used with SNPRelate (Zheng et al., 2012) to extract SNPs subsets of the 163,376 variants in GBS and reference samples above that were only from gene-rich or gene-poor regions. We then calculated Fst according to Hudson et al., (1992) separately for each chromosome using the R package KRIS v1.1.6 (Chaichoompu et al., 2018).

### Centromere identification

We identified centromeres using TRASHv1.2 (Wlodzimierz et al., 2023) with default settings. We also identified centromeric regions using CentIER (Xu et al., 2024) without including our gene annotations because of gff format incompatibilities. In general, overlap between the regions identified with CentIER and the satellites located by TRASH was incomplete. Since TRASH had successfully predicted a sequence also validated by FISH, we therefore focused on the TRASH results for downstream analyses. We compared sequence similarity of putative satellite sequences using ClustalW (Thompson et al., 1997).

With custom R scripts, we located the largest block of putative satellite repeats identified by TRASH that were within 100kb of each other in each genome. We then inspected the structures of these regions, buffered by 1Mb in either direction, when aligned to themselves using alignments generated with the nucmer command in MUMMER and plotted in R (Marçais et al., 2018). For all chromosomes in all genomes with the exception of chromosome six in MN106, the structure we observed was consistent with expectations). However, for chromosome six in MN106, the second largest contiguous block of satellite repeats showed a structure more consistent with a centromeric region.

### NLR identification

We identified NBS-LLR genes structurally in *Thlaspi arvense* using HMMER v. 3.4 (Eddy, 2011) with default settings by searching protein sequences against the Pfam NBS-LLR family (Pfam ID PF00931) raw hidden Markov model. For sequences with e-values above the HMMER default inclusion threshold (0.01), we then batch-searched the NCBI Conserved Domain Database using BLAST (https://www.ncbi.nlm.nih.gov/Structure/bwrpsb/bwrpsb.cgi), downloaded data as “full” (including both query and domain definitions), and for downstream analysis retained only the primary transcripts of proteins containing a complete NB-ARC family or superfamily domain.

Canonical NLRs contain a nucleotide binding domain and a leucine-rich repeat sequence, with variable specialized N- and C-terminal domains. We categorized pennycress NLR genes depending on the presence of three important N-terminal domains: Toll/Interleukin receptor (TIR), coiled-coil (CC), and RPW8 (RR8). With the same pipeline, we recovered 158 NLR genes from the *Arabidopsis thaliana* reference accession TAIR11, consistent with the lower end of literature estimates (Barragan and Weigel, 2021; Van de Weyer et al.,2019), which suggests that our methods are likely conservative but accurate. The distribution of core, dispensable, and unique orthogroups was similar regardless of whether orthogroups were present in arabidopsis (present: 47 core, 33 core single-copy, 11 variable, 3 unique; absent: 45 core, 35 core single-copy, 21 variable, 1 unique)

We used custom R scripts (github.com/kevinabird/pennypan) to extract phylogenetically hierarchical orthogroups containing NLRs and describe NLR clustering. Clusters were defined as at least two genes annotated as NLRs within no more than 50kb of each other. Syntenic blocks were identified using GENESPACE (see above). To describe synteny of NLR genes, we first extracted all blocks of synteny between the *Arabidopsis thaliana* TAIR11 genome and each of the seven pennycress genomes, calculated the total number of syntenic blocks for each pennycress genome relative to TAIR11, and determined how many syntenic blocks that overlapped an NLR cluster for each pennycress genome also overlapped an NLR cluster in TAIR11. To describe orthogroup synteny, we used the “.synHits” output from GENESPACE and queried whether the syntenic hits for each gene were in the same or different orthogroups, and performed a chi-squared test for each genome comparing NLR genes to non-NLR genes. Variability in NLR genes was calculated using per-site Shannon entropy according to the methods of Prigozhin and Krasileva (2021): NLR peptide sequences were aligned with mafft (https://pubmed.ncbi.nlm.nih.gov/12136088/) and per-site Shannon entropy was calculated using R scripts from Prigozhin et al., 2021. Alignments containing at least 10 peptides with a per-site entropy of at least 1.5 were considered “high-variability NLRs” (hvNLRs).

NLR sequence content saturation was calculated by identifying 31-mers shared by the seven accessions across NLR sequences and using these counts to generate growth curves with Pangrowth (Parmigiani et al., 2024b). Results from NLR analyses were compared against results for all genic regions and whole genome assemblies.

## Results

### Genome architecture and an updated reference for pennycress

While the existing ‘v2’ MN106 pennycress reference genome (Nunn et al., 2022) has served as a valuable resource, it also has a conspicuously large ‘bottom drawer’ of unanchored sequence that contains 75Mb (14.3% of the genome). There were also higher levels of duplicate sequence than may be expected, and optical map scaffolding can introduce false contig joins. As such, we opted to assemble a new ‘v4’ MN106reference with 88.35× PacBio HiFi (mean read length = 15,194bp), which was subsequently scaffolded with 91.4× OmniC and polished with 49.1× 2×150 Illumina reads. Combined, these methods produced one of the highest quality plant genome assemblies available to date.

Assembly required a total of five contig joins to form the seven chromosomes, there was no sequence in ‘bottom drawer’ contigs, all 14 chromosome termini were capped with long and likely complete putative telomere sequence, and four chromosomes were single-contig gapless (‘T2T’). Comparison between v2 and v4 assemblies revealed many major structural differences, including three large (> 1Mb in length) translocations (total = 24.7Mb) and 13 large inversions (total = 58.6Mb, Fig. 1A), none of which corresponded to contig break positions in v4 but many of which colocated with contig breaks in v2. These large likely artifactual inconsistencies cover > 18.5% of the seven v2 chromosomes. Generally, structural differences between the two genome versions are aggregated near putative centromeric positions, which is often the case in optical scaffolded assemblies, and also include an obvious 7.93Mb “deletion” where the centromere of chromosome one should reside.

**Figure 1.**
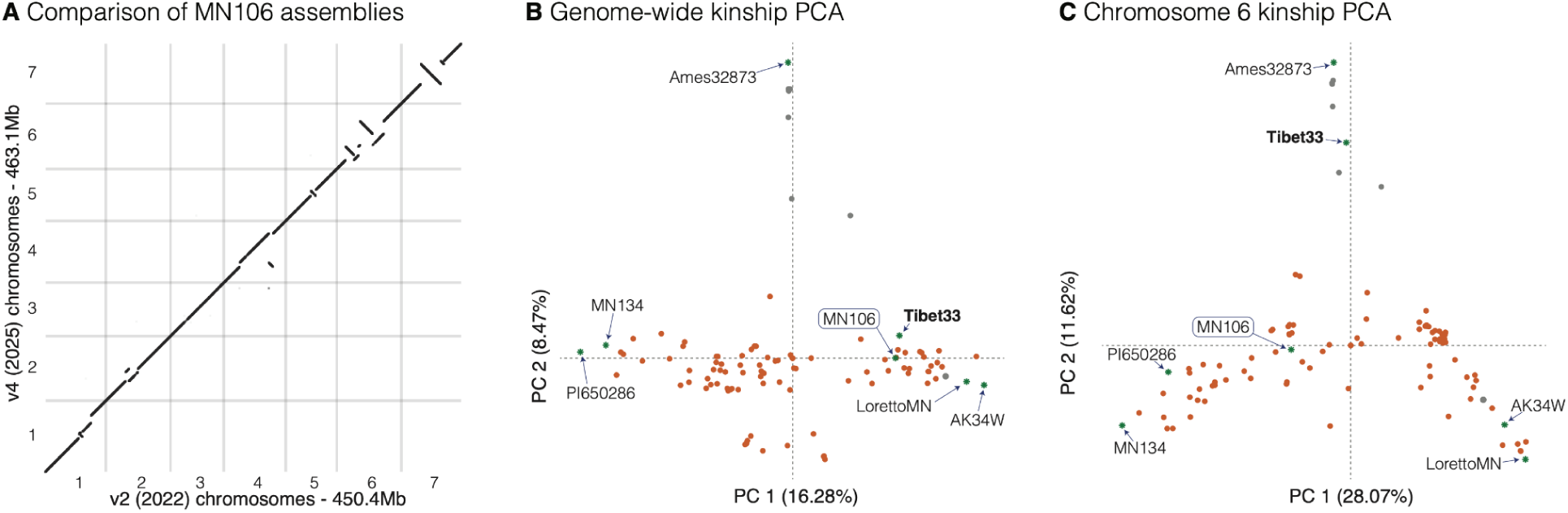
Genomic and sample diversity context for an updated MN106 reference genome. (**A**) minimap2 unique best (mapQ = 60) alignments of 145,577 non-overlapping 250bp ‘windowed’ v2 sequences aligned to the v4 MN106 assembly shows significant improvements in v4. For example, a large block of chromosome 1 is missing in v2 and inversions or translocations are present within all pericentromeres except that of chromosome 3. Principal component decomposition of kinship genomic relationship matrices calculated genome-wide (**B**) and using only chromosome 6 (**C**) show consistent patterns of relatedness among six of the seven reference genotypes. However, Tibet33 clusters tightly with the Armenian samples (dark grey points) in chromosome 6, potentially indicating a chromosome-scale introgression.

We complemented the v4 assembly with thorough annotations. Repeats were annotated using panEDTA in EDTA2 version 2.2.1 (Ou et al., 2019; 2024), telomeres by clustering of putative plant telomeric repeats, and centromeres by TRASH (Wlodzimierz et al., 2023) and CentIER (Xu et al., 2024, see below). Protein-coding genes were annotated with our ‘integrative’ approach that combines transcriptome, *ab initio* and homology support. Transcriptomes were built from 296M RNA-sequencing Illumina paired 2×150 reads and 8.7M full length transcript sequencing from a pooled PacBio ISO-seq library, *ab initio*, and homology support (see methods, Supplementary Data 2). While the v2 annotation was of fairly high quality (BUSCO = 98.7, 92% single-copy), our v4 represents a substantial improvement where protein coding genes capture 99.3% of embryophyte, and 100% of eukaryote, BUSCO genes.

Initial exploration of gene and repeat density revealed a structure typical of the *Brassicaceae* family and other angiosperms: gene content was mostly confined to dense regions in the chromosome ‘arms’ (20-23% genes, 19-26% repeats) while repeat sequences predominated in the ‘pericentromeric’ regions bounding the centromeres (0.9-1.2% genes, 85-87% repeats, Supplementary Fig. 1). However, unlike the closely related *Brassica rapa* and more distant *Arabidopsis* genomes that have small pericentromeres relative to chromosome arms, pennycress genomes are dominated by expansive pericentromeres that span 297-327Mb and make up 65-70% of the entire genome (Supplementary Fig. 1). This conspicuous structure allowed us to use a neural network approach to classify the genome sequences based on gene density, repeat density, and relative abundance of repeat types where sequence proximate to telomeres and centromeres served as training sets to bin genomic windows into ‘chromosome arms’ and ‘pericentromeres’ respectively (Supplementary Data 3). This classification clearly revealed a ‘two-speed’ genome structure, characterized by gene-rich chromosome arms collinear across genotypes (as gene-dense as the compact *Arabidopsis thaliana* genome), and large complex and possibly non-recombinant pericentromeric blocks (Supplementary Fig. 1).

### Abundant sequence and gene variation characterize the pennycress pangenome

To form a foundation for pennycress pangenomics, we selected six genotypes in addition to MN106 that spanned key breeding samples and corners of native genetic diversity, with origins from Alaska (‘AK34W’), Tibet (‘Tibet33’), Germany (‘PI650286’), and Armenia (‘Ames32873’) and two accessions that are foundational for North American breeding programs (‘MN134’, ‘LorettoMN’). Exploration of single nucleotide polymorphisms (SNPs) for 95 diverse genotyping-by-sequencing (GBS) pennycress lines and the polishing libraries of the seven pangenome reference genotypes confirmed that they capture the majority of diversity across the species (Fig. 1B). As previously noted (Frels et al., 2019; Nunn et al., 2022; Contreras-Garrido et al., 2024), samples from Armenia were particularly diverged. Within-chromosome clustering by genetic relatedness matrices also revealed some major discrepancies indicative of large-scale chromosomal introgressions. For example, ‘Tibet33’ is closely related to the Armenian group only on Chromosome 6, but is related to other genetic backgrounds on all other chromosomes (Fig. 1C), suggesting the effect of chromosome-scale introgression between deeply diverged subpopulations.

To dissect the genetic basis of divergence among these key genotypes, we assembled and annotated genomes for the six genotypes in an identical manner to, and achieved similar quality statistics as, MN106 (described above, Table 1). Our previous work (Lovell et al., 2018; 2021; Brůna et al., 2025) has shown that despite very high individual quality, independent annotations even with identical methods can artificially inflate gene presence-absence variation among pangenome members. We therefore applied a second ‘harmonization’ round of protein-coding gene model annotation where high-confidence peptides predicted across all genomes were projected onto each individual annotation. These homology-supported models replaced the original model if the resulting annotation had a higher score (see methods) or were added to the annotation if no independent gene model existed at the given locus. This approach produced between 27,470 and 28,765 high-accuracy gene models per accession (all Eukaryote BUSCO = 100%).

**Table 1.** Genome annotation, assembly and structure information across the seven pennycress genomes. *Two equally computationally-supported centromeric repeat arrays exist on Chr06 within genotype MN106; likely only one is recognized by the plant, but each is syntenic to the common centromere (left) or Armenia-only centromere (right).

|  | **MN106** | **Ames32873** | **AK34W** | **LorettoMN** | **MN134** | **PI650286** | **Tibet33** |
| --- | --- | --- | --- | --- | --- | --- | --- |
| Genome Size (Mb) | 463.07 | 460.15 | 456.03 | 455.92 | 461.65 | 466.53 | 456.74 |
| HiFi Coverage | 88.35x | 70.23x | 73.80x | 79.48x | 73.15x | 71.24x | 73.56x |
| Contig N50 (Mb) | 64.8 | 32.0 | 58.8 | 58.7 | 60.5 | 59.9 | 59.9 |
| N. Genes | 27,768 | 27,858 | 28,126 | 27,479 | 27,878 | 28,165 | 27,928 |
| Telomeres (n, Kb) | 14, 38.5 | 14, 31.6 | 14, 29.6 | 13, 36.4 | 14, 31.8 | 14, 29.6 | 12, 23.6 |
| Chromosome Arms (Mb, % genes) | 130.5, 23.0 | 139.4, 21.8 | 152.3, 20.6 | 153.8, 20.1 | 142.0, 21.6 | 158.5, 19.9 | 141.9, 21.7 |
| Pericentromeres (Mb, % repeats) | 327.6, 85.1 | 308.1, 85.0 | 299.2, 87.1 | 297.1, 87.2 | 314.5, 86.2 | 303.5, 87.8 | 308.4, 85.3 |
| Centromeres (n, Mb) | *8, 4.9 | 7, 12.6 | 7, 4.5 | 7, 4.9 | 7, 5.1 | 7, 4.5 | 7, 6.4 |

To assess pan-genomic variation in pennycress, we first created a pangenome graph with Minigraph-Cactus (v2.9.0; Hickey et al., 2023) using MN106 as the primary reference (Fig. 2A). On average, 95% of each assembly was retained in the final graph, which was 561.5 Mb in length and comprised 18,000,685 nodes and 24,228,161 edges. The core pan-genome, or sequences in the graph shared by all haplotypes, was 300.0 Mb (∼53%) in length. Of the remaining dispensable graph, 176.7 Mb (∼31%) of sequence was present in 2-6 haplotypes, and the remaining 84.8 Mb (∼15%) of sequence was unique to a single haplotype. Although our pangenome does not capture all possible sequence variation, the 1.254 decay parameter of the growth curve suggests that it captures the intended diversity at the base level (Parmigiani et al., 2024b).

**Figure 2.**
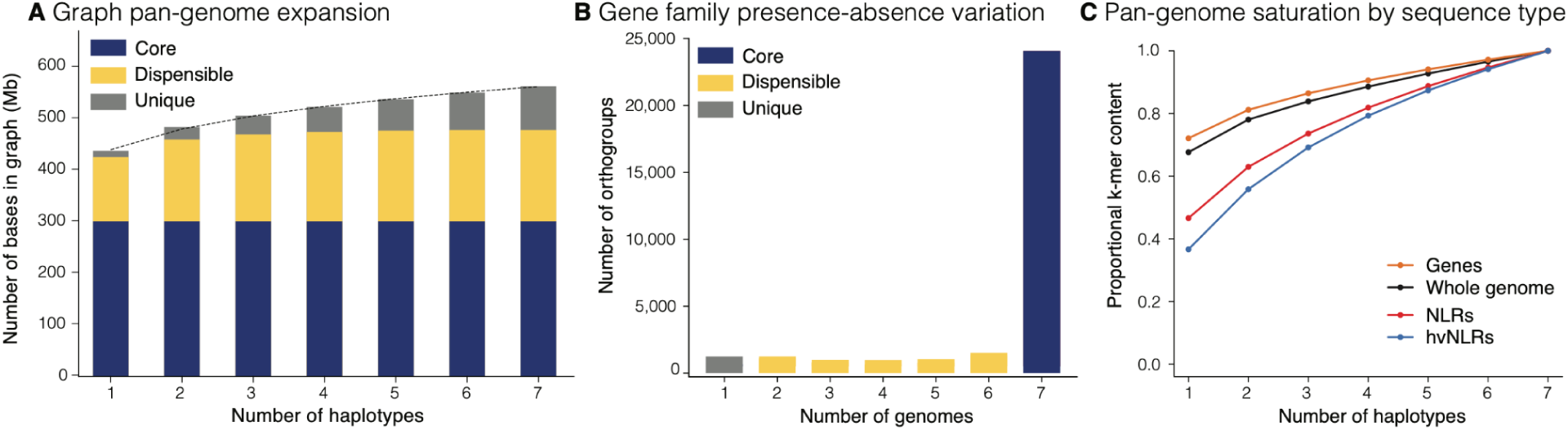
Pan-genome variation in pennycress. **A** The amount of sequence in the genome graph that is shared by different numbers of haplotypes (assemblies). This figure only represents sequence in the final graph and does not contain sequences ‘clipped out.’ **B** Breakdown of core, dispensable, and unique orthogroups among our seven genomes. **C** Growth curves describing patterns of 31-mer sharing across assemblies; a steeper curve indicates a smaller degree of k-mer sharing among assemblies for that sequence class.

We next queried the extent of presence-absence variation (PAV) at the gene family level using 27,226 hierarchical orthogroups identified by the phylogenetic orthology inference method Orthofinder2 (Emms and Kelly, 2019) with *Arabidopsis thaliana* as an outgroup (Fig. 2B; Supplementary Data 4). Although only 53% of the total DNA sequence was identified as core, 80.3% of the orthogroups (24,077) were core, and 91% of core orthogroups (20,985) were uniformly single-copy across all seven assemblies. For the remaining orthogroups, 4,666 (16%) were present in 2-6 genomes, and 1,242 (4%) were unique to a single genome. The discrepancy between orthogroup PAV and variable sequence in the genome graph is substantially larger than reported in other recent pangenomes, perhaps indicating that our efforts to harmonize annotations across genomes effectively reduced artifactual PAV (Brůna et al., 2025).

We also applied a kmer-based approach to describe the amount of core and dispensable sequence (Fig. 2C), and found that sequence sharing in genic regions exhibits a similar pattern to the orthogroup-based analysis. However, it should be noted that known highly variable genes like NLRs (see below) do show substantially more sequence PAV than the full genome and genic sequences. The disparity between low presence-absence variation in the highly syntenic genic regions and higher DNA sequence variation reflects the uneven distribution of structural variation across the ‘two-speed’ genome structure. Although the pangenome may not capture all sequence variation in pennycress, the high sequence sharing in genic regions indicates that it does contain the vast majority of important functional variation relevant to crop improvement.

### Extensive structural variation in pericentromeres and two-speed evolution across pennycress genomes

Genotypes sampled within our pennycress pangenome are characterized by highly syntenic chromosome arms, but extensive structural rearrangements (SVs) throughout gene-poor pericentromeres (Fig. 3). Similar to our SNP-based clustering, SVs were most common between the deeply diverged Armenian Ames32873 genome and the others. For example, SyRI classifications of alignments revealed that the five genomes excluding Tibet33 (see below) were syntenic to MN106 over more than 330Mb (75%), but in Ames32873 only 122Mb of sequence (24%) was syntenic to MN106 (Supplementary Figure 2A-B), mainly attributable to un-alignable sequence (classified as either ‘highly-diverged regions’ (HDRs) or unaligned regions; Goel et al., 2019). These regions are most likely complex nested genomic rearrangements (Fig. 3). In contrast to consistent genome wide divergence relative to the Armenian genome, the genome-wide synteny tracking revealed high structural similarity between Ames32873 and Tibet33 for chromosome six (Fig. 3) but similar structures between Tibet33 and the other genomes on the other chromosomes. These SVs are consistent with SNP-based evidence for a single chromosome-scale introgression (Fig. 1C).

**Figure 3.**
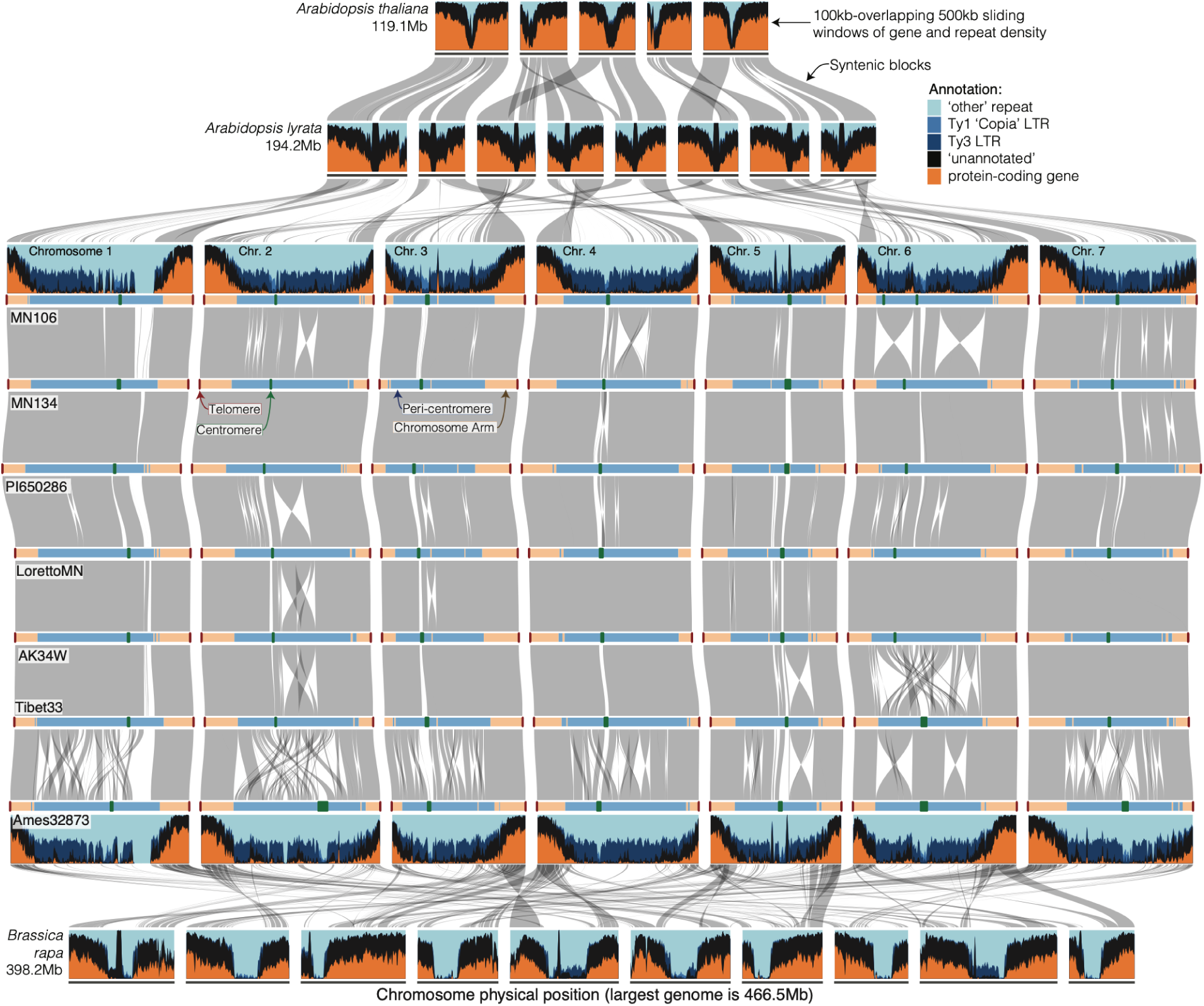
Macrosynteny and genome structure across the Brassicaceae. Horizontal blue/black/orange bands represent the chromosomes of *Arabidopsis thaliana*, *A. lyrata*, MN106, and *Brassica rapa* (top to bottom). Chromosomes are ordered by their number from left to right. Colors represent genomic content binned hierarchically in sliding windows (400kb-overlapping 500kb) as follow: (1) within a gene annotation (including intron and UTR, orange), (2) within EDTA-annotated repeats categorized as Ty3, (3) Ty1 (copia), (4) within another repeat category, or (5) un-annotated. Grey bands are sequence-based syntenic blocks between each pair of genomes. Pennycress and *B. rapa* are phylogenetically proximate (both in Brassicodae supertribe), but have reduced synteny in part because of genome reshuffling in *B. rapa* following a whole-genome triplication event. The seven pennycress genome assemblies (horizontal bars) are binned into TRASH-defined centromeres (orange), pericentromeres (dark blue), chromosome arms (light blue) and telomeres (dark red). The colors along the chromosome segments scale physically with the size of the bin, except that centromeres and telomeres have a 1pt buffer to make it easier to see these typically small regions. Each genome is connected to its neighbor by grey polygons that represent sequence-based syntenic blocks. Plots, genomic bins, and syntenic blocks were built with DEEPSPACE (github.com/jtlovell/DEEPSPACE).

Overall, the massive level of complex structural variation in the pericentromeric regions stands in stark contrast to the largely conserved genomic content in the telomeric arms, and illustrate how some regions of the genome are more permissive of structural variation in pennycress. Structural variation can create complications in meiosis that limit gene flow within and between species. This has been widely noted for inversions (e.g. Rieseberg et al., 1999; Noor et al., 2001; Rieseberg, 2001; Kirkpatrick and Barton, 2006; Navarro and Barton, 2006; Noor et al., 2006; Coughlan and Willis, 2016), and is increasingly appreciated for other forms of rearrangements, such as dysploid rearrangements (Beaudry et al., 2022) and centromere repositioning (Steckenborn and Marques 2025). We therefore sought to assess if large rearrangements within the centromeric and gene poor regions of the Armenian accession affect gene flow in pennycress. For this, we calculated pairwise *F_ST_* (Hudson et al., 1992) from our SNP data between Armenian and non-Armenian accessions to assess patterns of differentiation between gene-rich and gene-poor regions. Across all chromosomes, we found *F_ST_* to be higher in the gene-poor, more rearranged pericentromeric regions (median Fst pericentromere = 0.54, chromosome arms = 0.48, Wilcoxon rank sum test p < 0.05; Supplementary Figure 3). Importantly, non-genic regions are often under weaker evolutionary constraint and (peri)centromeric regions generally have lower recombination rates, both of which are expected to increase *F_ST_*. Thus, pennycress inversions, particularly in pericentromeric regions may decrease the potential for recombination and act as a barrier to gene flow.

### Dynamic evolution of centromeric satellite repeats

Synteny analysis revealed extensive rearrangement in centromeric regions, especially relative to the Armenian accession. However, it remained unclear whether this variation affected the centromeres, which are essential for meiosis and chromosome pairing, or merely the highly repetitive pericentromeric regions more likely to experience relaxed selection. We sought to better characterize the contents of repeat-rich pericentromeric and centromeric regions by identifying known pennycress centromeric satellite repeats and regions using TRASH (Wlodzimierz et al., 2023) and CentIER (Xu et al., 2024). TRASH, which discovers repeats from regions with high kmer repetitiveness, identified five putative satellites ranging in size from 73bp to 648bp that constituted 3.1%-5.6% of total genome content and were concentrated in regions containing 50%-80% satellite sequence per 1kb window (Supplementary Data 5; Supplementary Fig. 3). The most abundant satellite (1.7-3.1% of total content), a 166-bp sequence, occurred on all chromosomes in all accessions and was identical to a satellite previously identified from a pennycress accession from Brno, Czech Republic using fluorescent in-situ hybridization (FISH) of sequences derived from BAC clones (Bayat et al., 2021). Its abundance and wide distribution suggest it may be ancestral in pennycress. Putative centromeric regions identified by CentIER showed only imperfect overlap with regions enriched for this FISH-validated satellite. We therefore designated the approximate location of the centromere as the largest contiguous cluster of repeats within 100kb of each other of either the 166bp or the 73bp satellite sequence (which was abundant on chromosome five in all accessions).

Both centromere repeat structure and position varied across accessions. In particular, the Armenian line (Ames32873) had extensive rearrangements in all centromeric regions and more total putative centromeric satellite sequence (5.6% of total sequence, compared to 3.1%-4.3% in other accessions; Fig. 4), and when inferred centromeric regions were aligned to themselves, variation in repeat structure was apparent, both within and between genomes. For example, the bounding 100kb of alignable sequence adjacent to putative centromeres is completely syntenic between Ames32873 and MN106 only on chromosome 4. Conversely, the syntenic position and bounding sequence identity is completely different between centromeres of chromosomes 1 and 6. These results strongly suggest ongoing evolution of centromere architecture and genomic positioning (Fig. 4), which may underlie incompatibilities that could impact breeding efficiency between deeply diverged gene pools.

**Figure 4.**
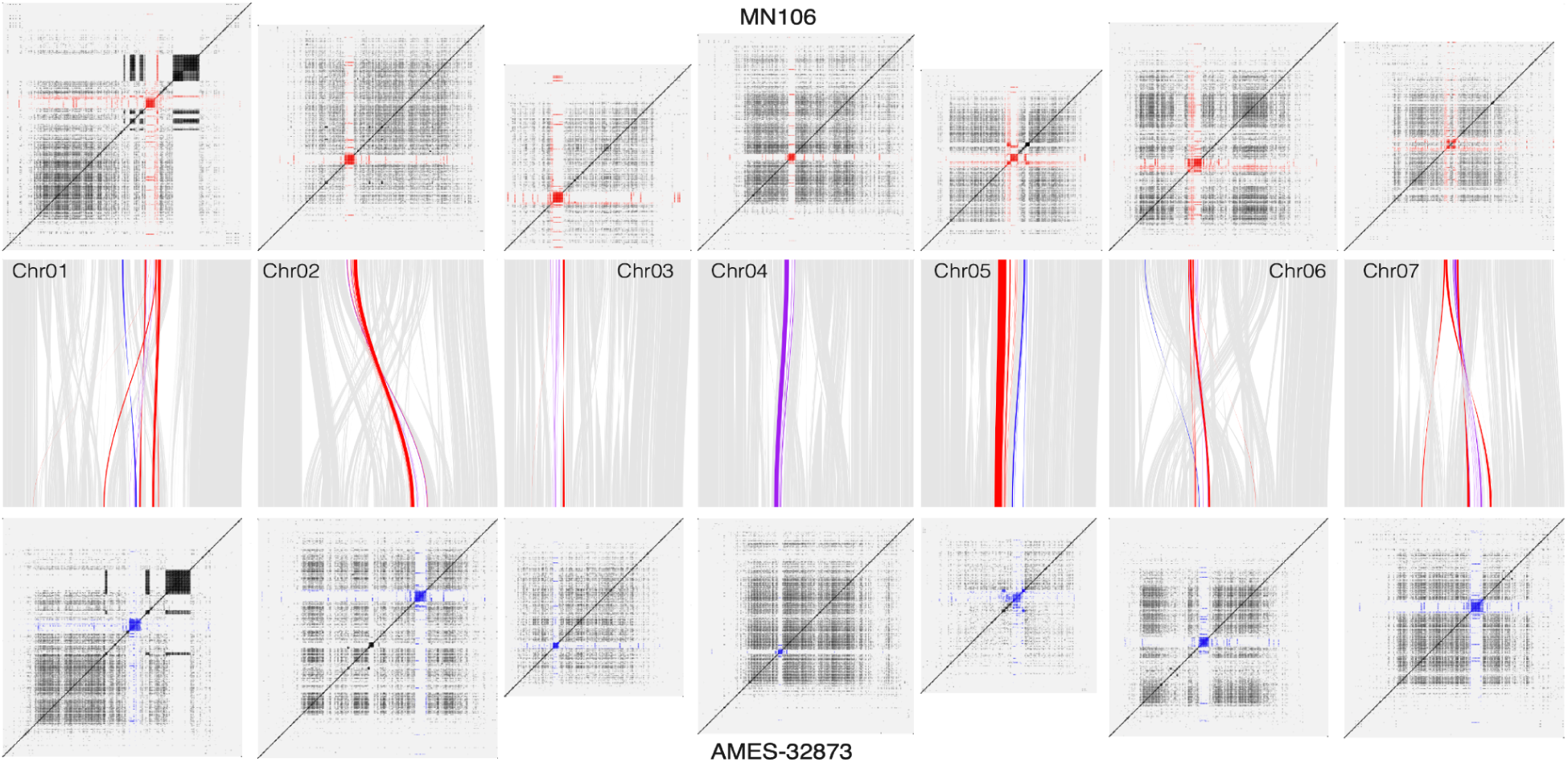
Synteny between centromeric satellite repeat regions in pennycress ‘MN106’ (top, red) and pennycress ‘Ames 32873’ (bottom, blue). Chromosome panels (top and bottom): self sequence mapping 24-mers (dark regions indicate repetitive sequence). Central panel: gray ribbons indicate background sequence synteny, red and blue ribbons indicate synteny for satellite sequences in the largest contiguous block of satellite repeats on a chromosome.

### Rapid evolution of pathogen resistance genes in the dispensable genome

Although the pangenome growth curves suggest that the gene-dense portion of the genome is comparatively conserved (Fig. 2C), the presence of dispensable and unique genes reflects variation in content and potentially in evolutionary rates. To characterize this variation, we performed gene-ontology enrichment (GO) analysis on Arabidopsis genes in orthogroups annotated as either dispensable or unique in pennycress. The dispensable genome (‘shell’) was enriched for genes annotated as involved in cell-cell signaling (p = 6.6 x 10^-12^, Fisher’s exact test), and unique genes (‘private’) were enriched for defense response (p = 9.4 x 10^-6^, Fisher’s exact test). The enrichment for defense related genes in highly variable portions of the genome suggests that pangenomic analyses could benefit pest and pathogen resistance, a central breeding target in pennycress (Basnet and Ellison, 2023). We therefore focused on annotating and describing the evolution of a major class of disease-resistance genes.

NOD-like receptor (NLR) genes are a major group of disease-resistance genes in plants, known to be enriched for functionally important copy number and presence absence variation in many species (Barragan and Weigel, 2021; Lovell et al., 2021; Chou et al., 2023). We applied our pangenomic approach to characterize NLR diversity and aid pennycress breeding efforts to reduce vulnerability to diseases and pests, including soybean cyst nematode. We first structurally identified between 131 and 148 NLRs per accession with three canonical N-terminal domains (Supplementary Table 1; Supplementary Data 6). Analyses using kmer and orthogroup level variation demonstrate higher variability in NLR genes: NLRs have a substantially steeper growth curve than other genic sequence (Fig. 2C), indicating more PAV and NLR enrichment in the dispensable genome. Similarly, across the 132 phylogenetically hierarchical NLR orthogroups we identified, 52% were private to pennycress (not found in Arabidopsis) and NLR genes were slightly less likely to be included in core orthogroups (81% vs 85%, p=0.2298, Fisher’s exact test), somewhat more frequent in orthogroups present in 2-6 genomes (15% vs 12%, p=0.1339, Fisher’s exact test), and significantly less common in core orthogroups that are single-copy in all accessions (58% vs 77%, p = 6.47 x 10^-7^, Fisher’s exact test).

Although it is known that NLR genes tend to group into physical clusters (van Wersh and Li, 2019), less is known about variation in the position and content of NLR clusters within and between species. Characterizing higher-level NLR cluster evolution of this kind may help both identify candidate resistance loci in emerging crop systems and shed light on the evolutionary dynamics of these complex genomic loci. We leveraged our highly contiguous pangenome assemblies to track NLR clustering and cluster synteny within pennycress, identifying between one and 14 clusters of between two and 11 NLRs per chromosome (Supplementary Table 1). Although NLR clusters are rare in these genomes overall (<16% of sequence), the vast majority (>90%) of pennycress NLR clusters were syntenic to regions that also contained NLR clusters in Arabidopsis (Supplementary Table 2). While the physical position of NLR clusters is highly syntenic between pennycress and Arabidopsis, NLR orthogroups in syntenic clusters were similar within pennycress but differed between species, suggesting that rapid turnover erodes 1:1 synteny (Fig. 5). Consistent with this, NLR genes located within a cluster shared across species (pennycress NLRs with syntenic BLAST hits to Arabidopsis NLRs) were significantly less likely to be in the same orthogroup than other genes genome-wide (chi-sq with 1 df, all p<0.0001). Thus the macrosyntenic conservation of NLR clusters between Arabidopsis and pennycress does not reflect conservation of NLR gene content within these clusters.

**Figure 5.**
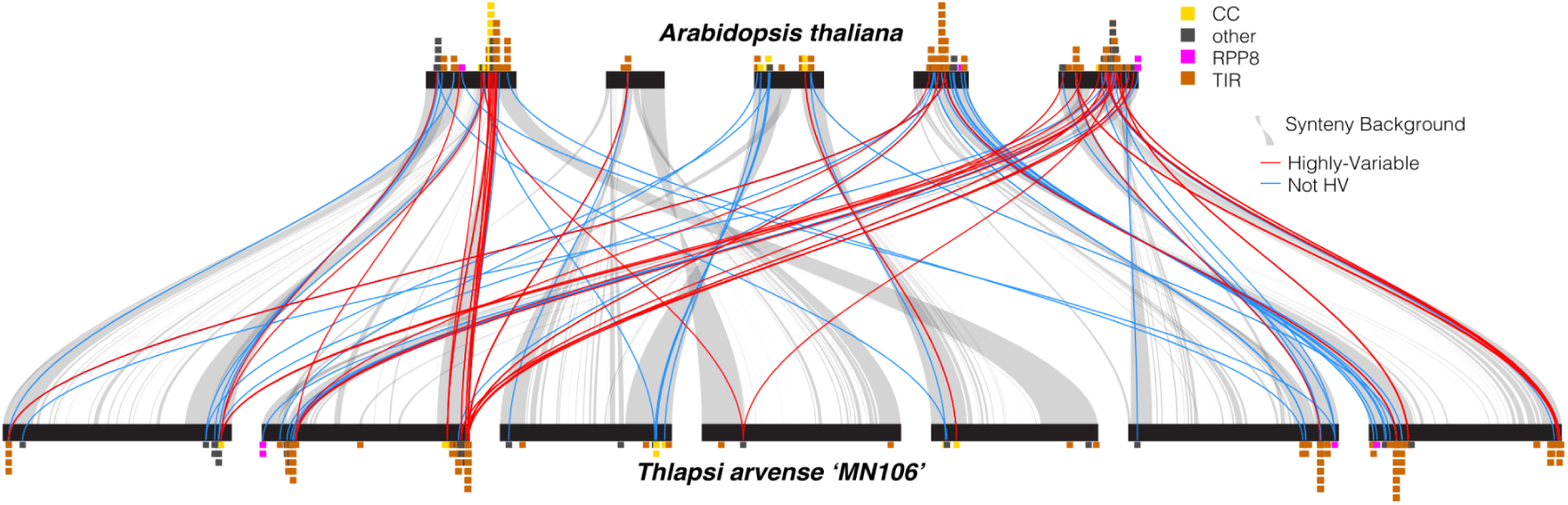
Synteny between locations of NLR clusters in Arabidopsis (top) and pennycress ‘MN106’ (bottom). Stacked barplots represent NLR clusters (one block per gene), colored by N-terminal domain. Gray ribbons indicate synteny. Red ribbons indicate synteny between highly variable NLR genes (hvNLRs), and blue ribbons indicate synteny between non-hvNLRs.

The variation in content of syntenic NLR clusters also suggests that distinct sets of NLRs are diversifying between species, likely reflecting their unique ecologies and evolutionary pressures. We characterized NLR variability (evolutionary rates) in pennycress by calculating per-site Shannon entropy for NLR gene-containing orthogroups and related this to a previously identified set of highly variable NLR genes in Arabidopsis (Prigozhin and Krasileva 2021). We identified 9% of NLR gene-containing orthogroups (12/132), representing a total of 244 genes, as having sufficient peptide variability to count as high-variability NLRs (hvNLRs). All eight orthogroups containing Arabidopsis hvNLRs were present in pennycress, and six of these varied in gene number. However, only seven of the 12 orthogroups containing pennycress hvNLRs were present in Arabidopsis, and only three were high-variability in both species. Although orthogroups containing Arabidopsis hvNLRs were more likely to contain pennycress hvNLRs, the difference was not statistically significant (chi-sq with 1 df, p = 0.073), suggesting that different NLR orthogroups evolve rapidly in pennycress compared to arabidopsis. Taken together, these analyses of NLR diversity and synteny suggests that while NLR clusters are largely positionally conserved across species, species vary greatly in their NLR content within clusters and the identity of expanding NLR orthogroups and fast-evolving hvNLRs differs between species (Fig 5). This both identifies specific high-variability NLR clusters within pennycress that could be a focus for breeding efforts to shift biotic resistance and offers insights into the processes governing NLR evolution and diversification in the Brassicaceae. This class of important genes also illustrates the complexity of the ‘two-speed’ genome compartmentalization: rapid evolution occurs in syntenic, gene-dense regions, but even with high gene turnover and extensive presence-absence variation, synteny is maintained.

## Discussion

In this study we leveraged the latest long-read sequencing technology to make seven exceptionally high-quality genomes representing the genetic and geographic diversity of the emerging oilseed crop pennycress (*Thlaspi arvense*). All seven genomes showed a ‘two-speed’ genome compartmentalization pattern of small extremely gene-dense regions at the ends of chromosomes and large pericentromeric regions that are overwhelmingly comprised of transposable elements (TEs) and other repetitive sequence. Interestingly, these large pericentromeric rearrangements resulted in minimal variation in the size of the genomes, suggesting they operated within a narrow size range of DNA Kbp. In the genic regions, our pangenomic analyses revealed that the vast majority of orthogroups and sequences were shared across all genomes. We uncovered a moderate amount of structural variation in the form of large inversions, and interchromosomal translocations were entirely absent. These results show that the relative paucity of SNP variation in the species (e.g. Frels et al., 2019) is mirrored by generally low levels of pangenomic variation for most genic content in the genome. We did, however, identify extensive variation in two aspects of the pennycress genome: the structure of the centromeres, and the complement of immune defense NOD-like receptors (NLR) genes. Because both structural variation in highly repetitive regions such as centromeres and presence-absence variation in gene complement are hard to characterize using reference-based approaches, our discoveries highlight the value of pangenome approaches.

### Structural and centromeric diversity

Our pangenomic analysis revealed extensive genomic rearrangement (∼25% sequence syntenic) between the Armenian sub-population and our six other genomes that far exceeds what has been observed in pangenomes of other species in the Brassicaceae including *Arabidopsis thaliana* (70%-95% syntenic; Lian et al., 2024), *Brassica napus* (74-86% syntenic; Song et al., 2020), and *Camelina sativa* (∼94% syntenic; Bird et al., 2025). It also exceeds other recent pangenomes like tetraploid potato (56% syntenic between 2 haplotypes; Sun et al., 2025) and *Eucalyptus* (50-53% syntenic between the most distinct subsets of species; Ferguson et al., 2024), despite pennycress being a relatively young diploid species (Esmailbegi et al., 2017). Recent reports in Mimulus are qualitatively similar, where large blocks of unalignable sequence in pericentromeric regions resulted in ∼75% synteny within a *M. guttatus* population and ∼43%-54% synteny between different *Mimulus* species, but without the centromeric shifts observed in this study (Lovell et al., 2025). While this magnitude of genomic divergence raises questions about the proper taxonomic status of the pennycress Armenian population, there are no known reports of unsuccessful or onerous crosses between Armenian and non-Armenian accessions (Frels et al., 2019) and our results suggest at least one chromosome-scale introgression between the Armenian Ames32873 line and Tibet33. Similar signs of unidirectional gene flow from Armenian populations were seen in a recent large-scale population genomic survey of pennycress (Wu et al., 2025). Previous studies established the Armenian subpopulation as genetically diverged from other populations based on SNP data (Frels et al., 2019; Nunn et al., 2022; Contreras-Garrido et al., 2024; X. Wu et al., 2025) and, on this basis, proposed it as a breeding source to expand genetic diversity in the species (Frels et al., 2019). However, because the diversity is primarily in non-functional pericentromeric regions and the Armenian genome did not stand out as distinct in its PAV or NLR repertoire, its value as a source of genetic diversity appears limited. Instead, this pennycress system presents an exceptional model to investigate the causes and evolutionary consequences of widespread genomic rearrangements. It also shows the importance of diverse sampling across geographical and genetic distances for uncovering structural variation, even in species with apparently low SNP diversity.

This previously unappreciated structural variation also offers insight into ongoing debates about the distribution of genetic variation across chromosomes and the forces shaping these distributions. Across taxa, repeated observations have variously found higher and lower variation in pericentromeric regions. Possible explanations for these patterns have included gene density, recombination, selection, hitchhiking, domestication, and mating system (Aguadé et al., 1989, Kawabe et al., 2008, Flowers et al., 2012, Chen et al., 2022); here, we provide evidence that different forms of variation may predominate in different parts of the chromosome, with structural variation being relatively more important in less gene-dense regions. While our results do not identify the underlying cause of the extensive genomic rearrangement, many likely candidates exist. Transposable elements and other repetitive sequences are known to promote genomic instability, structural variation, and rearrangements in plant and animal genomes (Bennetzen, 2005; Balachandran et al., 2022). Rearrangements in pennycress are concentrated in the repeat-rich pericentromeric and centromeric regions in which TE activity is known to be concentrated (Contreras-Garrido et al., 2024). Centromere variants are also associated with meiotic drive in plants (Finseth, 2023), and the presence and abundance of drivers may vary spatially (Lindholm et al., 2016). Comparisons between the Armenian and non-Armernian populations may also shed light on how genomic rearrangements contribute to speciation or limit the efficacy of back-crosses through reduced recombination that prevents breaking linkage disequilibrium (LD) between beneficial and deleterious variants.

Because we found evidence of extensive rearrangement in the middle of the pennycress chromosomes, we used our highly contiguous independently assembled genomes for an in-depth query into the dynamics of complex centromeric regions. Using both previously validated and new computationally identified centromeric satellites, we estimated centromere position and structure for each chromosome across all seven assemblies, revealing that all seven centromeres were structurally rearranged and relocated in the Armenian genome, and that the repeat structures and locations also varied across non-Armenian samples. Previous work in the Thlaspideae tribe has found numerous changes in centromere position and structure between genera and species, and identifies chromosome six in particular as a rearrangement-prone driver of diversification in these species (Bayat et al., 2021). Our results demonstrate that these processes also occur at the intraspecific level, at least in pennycress. While several recent high-quality genomic studies have noted centromere movement within species (e.g. grape, maize, octoploid strawberry, soybean, and hexaploid wheat: Hufford et al., 2021; Y. Liu et al., 2023; Zhao et al., 2023; L. Guo et al., 2025; X. Jin et al., 2025), these mostly involve a subset of chromosomes and occur through canonical epigenomic centromere repositioning without changes to the underlying DNA sequence. The observations here of genomic rearrangements relocating every centromere between subpopulations are, to our knowledge, unprecedented. Pennycress presents a unique case study of genome instability and population divergence. We are unable at present to tell if centromeric movement occurred before the genomic rearrangements as part of canonical centromere repositioning or as a consequence of the rearrangements, or how the appearance of possible novel satellite repeats relates to this process. Future work is needed to validate centromere location with CENH profiling and formally test whether genomic rearrangements were the cause or consequence of centromere movement to shed light on possible evolutionary causes.

### Evolutionary history of NLR Clusters and implications for resistance breeding

Immune function genes evolve rapidly in many taxa (Pancer and Cooper, 2006). In particular the NOD-like receptor (NLR) genes in plants have long been known to be highly diverse (Baggs et al., 2017), and vary widely in complement and structure in other Brassicaceae species (Van de Weyer et al., 2019; Jiao and Schneeberger, 2020; Zhang et al., 2021). Because pennycress’s susceptibility to pathogens currently limits its use in crop rotations (Basnet and Ellison, 2024), we characterized NLR diversity in our pangenome to accelerate breeding for disease resistance (Barragan and Weigel, 2021). Although the pennycress pangenome generally showed strong conservation of gene order and only modest PAV, variation was unevenly distributed across the genome. As in other plants, pennycress NLR genes exhibit a more ‘open’ pangenome than other genes, showing evidence of high variability and rapid evolution.

As more sequenced genomes become available, species-specific ‘panNLRomes’ have revealed extensive variation in evolutionary rates and diversity as well as in structure, function, and genomic clustering (Van de Weyer et al., 2019; Lee and Chae, 2020; Prigozhin and Krasileva, 2021). Recent cross-species integrations also suggest that NLRs may maintain some level of synteny even across extremely long evolutionary timescales (B. Guo et al., 2025), and reveal the breadth of functional expansion in ancient, positionally conserved clusters (VanGessel et al., 2025). With the capacity to resolve and position tandem arrays across disparate genomes, the field moves toward a more complete general picture of NLR variability at the sequence, content, and position scales and the relationship between variation at different scales. Indeed, the NLRs we identified in pennycress grouped genetically into many orthogroups and were physically concentrated in clusters distributed across the genome. Our comparative genomic investigation of pennycress and Arabidopsis NLRs offers clear evidence for NLR cluster positional conservation across species. However, rather than reflecting shared gene content, the pennycress NLR clusters we identified included several orthogroups absent from Arabidopsis and variable orthogroup content across species. Peptide entropy estimates of pennycress NLR orthogroups (Prigozhin and Krasileva, 2021) further showed that high-variability orthogroups in pennycress were largely different than the high-variability orthogroups in Arabidopsis.

Taken together, these results support an evolutionary model where NLR cluster locations are broadly conserved but their component genes are frequently relocated and evolve rapidly, leading to distinct trajectories in cluster content and diversity across species. This erosion of 1:1 gene synteny within positionally conserved clusters poses unique challenges to translational biology as it means winnowing down candidate genes within a cluster based on orthology may be ineffective or misleading. Functional characterization of NLR variation in pennycress could complement our genomics approach to enhance breeding.

## Conclusions

Genomes are not homogeneous, but rather vary in a myriad of features including local recombination rate, gene and transposable element content, extent of heterochromatinization, and effective population size and efficacy of selection. The distributions of these features structure the degree and kind of variation in different genomic regions. Our results revealed pennycress as an extreme case of a compartmentalized genome structure resulting in two ‘speeds’ of pangenomic evolution: largely stable and conserved gene-rich chromosome arms, and gene-poor pericentromeric regions with extensive structural variation. In a pangenomic context, our findings highlight both NLRs and centromeres as forms of genomic content that undergo substantial rearrangement and restructuring that would not be detectable using reference-based approaches. In both cases, the functional content of the region remains the same but the exact gene or repeat content displays repositioning and presence-absence variation. Plant pangenomes are now allowing us to characterize these forms of previously elusive variation, with exciting implications for our understanding of their evolution.

## Acknowledgements

The work (proposal: Award DOI 10.46936/10.25585/60001293) conducted by the U.S. Department of Energy Joint Genome Institute (https://ror.org/04xm1d337), a DOE Office of Science User Facility, is supported by the Office of Science of the U.S. Department of Energy operated under Contract No. DE-AC02-05CH11231. This work was supported by National Science Foundation Division of Integrative Organismal Systems grant 2208944 to KAB, United States Department of Agriculture National Institute of Food and Agriculture grant 2019-05709 and National Science Foundation Division of Molecular and Cell Biology grant 1906486 to DJK, and National Science Foundation Division of Integrative Organismal Systems grant 2029959 and Department of Energy Office of Biological and Environmental Research grant DE-SC0022987 to PPE

## Data Availability

Reference genome and annotation files for Thlaspi arvense of AK34W (v 1.1), Ames32873 (v1.1), LorettoMN (v 1.1), MN106 (v4.1), MN134 (v1.1), PI650286 (v1.1), and Tibet 33 (v1.1) genomes are available at https://phytozome-next.jgi.doe.gov/pennypan/. All raw genomic sequence reads have been deposited in the NCBI SRA database under BioProject accessions PRJNA1321803,PRJNA1321804,PRJNA1321814, PRJNA1321815,PRJNA1321823 PRJNA1321824,PRJNA1321826. All raw transcriptomic sequences used for annotation have been deposited in the NCBI SRA database, their BioProject accessions and other metadata are listed in Supplementary Data 2. Raw data for GBS resequencing can be found in the NCBI SRA database under BioProject Accession PRJNA1315838

## Supplementary material

**Supplementary Data 1 | Genome sequencing and assembly statistics.**

Summary of genomic sequence coverage, read lengths, and genome assembly contiguity for all genotypes used in this study.

**Supplementary Data 2 | Metadata for RNA- and ISO-seq libraries used in protein-coding gene annotations**. Genotype (and also genome) identifier, internal unique library ID, total number of reads, total number of reads mapped to the reference genome, sequencing type, and NCBI BioProject Accession are reported for each sample.

**Supplementary Data 3 | Genome classifications**. Each genome was split into 5kb-overlapping windows and classified into three bins (subtelo aka chromosome arm, pericentromere, centromeres) by the repeat and gene content therein. Coordinates of blocks of adjacent bins are reported for each genome and chromosome.

**Supplementary Data 4 | Gene family identity**. GENESPACE-formatted global and synteny-constrained OrthoFinder results.

**Supplementary Data 5 | Centromere hits**. Repetitive regions identified by TRASH from all seven genomes for all five satellite sequences with cluster membership and cluster locations.

**Supplementary Data 6 | NLR genes**. Concatenated GFF containing N-terminal domains, gene locations, cluster membership, high-variability status, and orthogroup IDs for all NLR genes identified across all genomes.

