## Supplementary Table 1 for "Structure and sequence evolution in the pennycress (*Thlaspi arvense*) pangenome"

Online supplementary material for: structure and sequence evolution in the pennycress (*Thlaspi arvense*) pangenome

**Supplementary Figures	1**

Supplementary Figure 1 | Gene and repeat density	1

Supplementary Figure 2 | Identity of alignable and unaligned sequence relative to MN106	2

Supplementary Figure 3 | Population differentiation by genomic class	2

Supplementary Figure 4 | Position and structure of centromeric repeats	3

**Supplementary Tables	4**

Supplementary Table 1 | NLR gene and cluster counts by accession.	4

Supplementary Table 2 | NLR cluster syntenic block overlap.	4

#### Supplementary Figures

##### Supplementary Figure 1 | Gene and repeat density

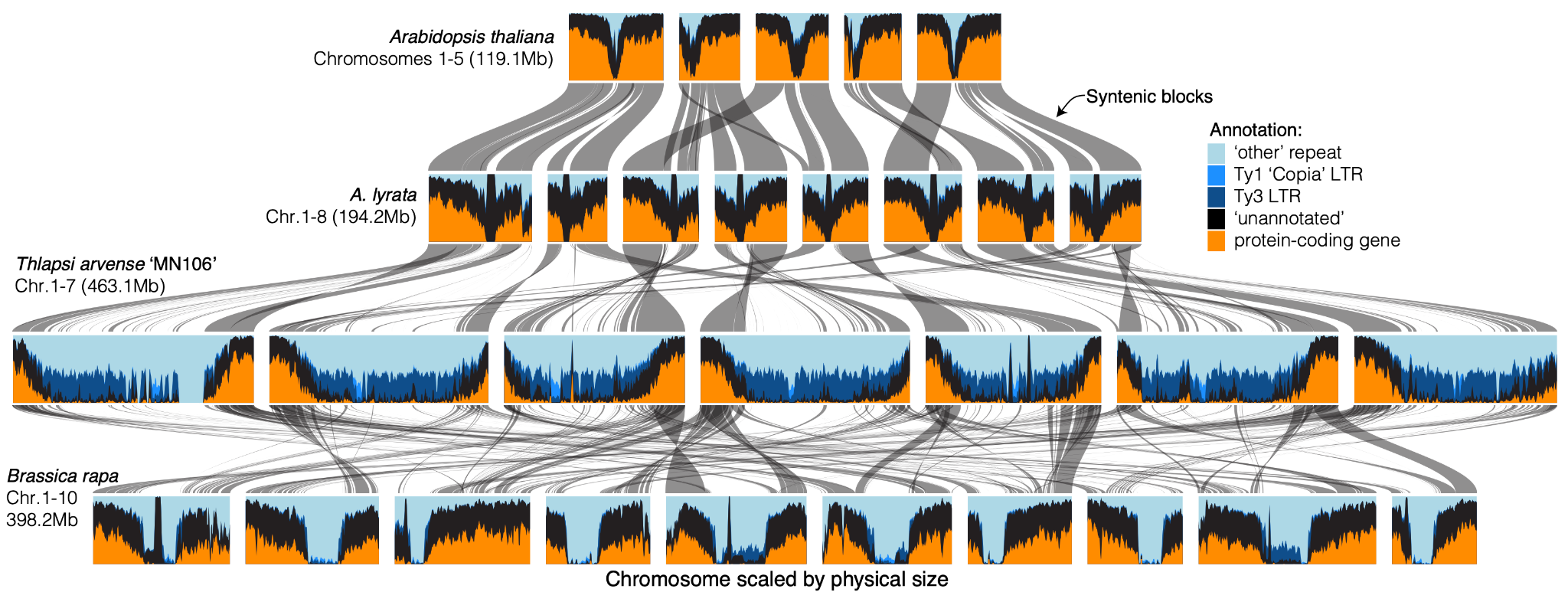

**Supplementary Figure 1 | Gene and repeat density.** Synteny map and sliding windows following Fig. 3. See descriptions therein for details.

##### Supplementary Figure 2 | Identity of alignable and unaligned sequence relative to MN106

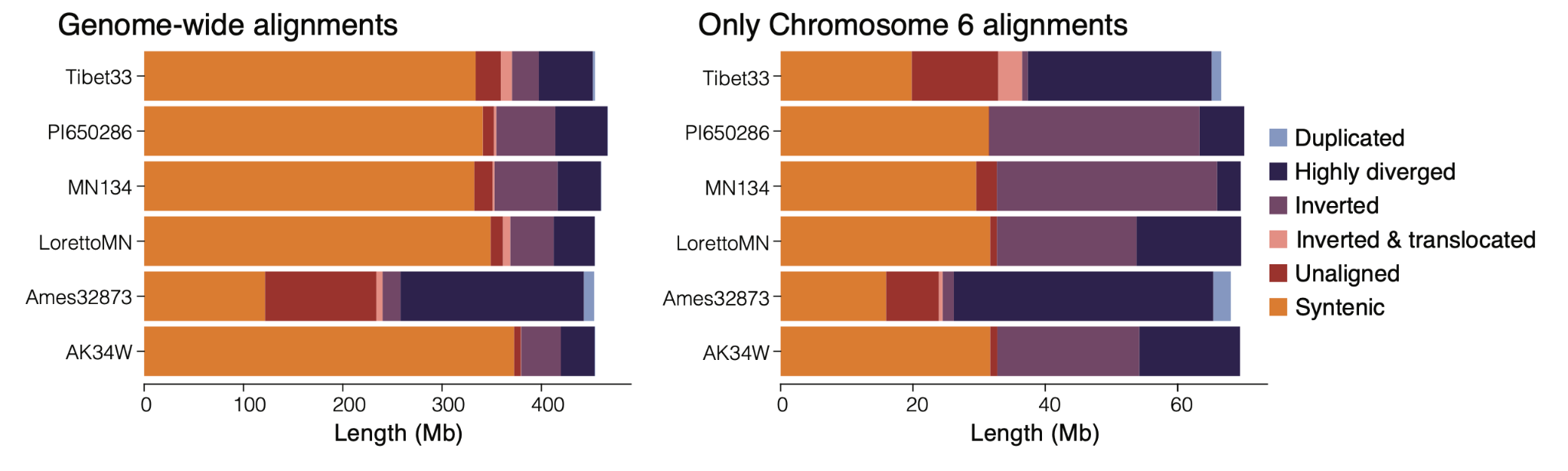

**Supplementary Figure 2 | Identity of alignable and unaligned sequence relative to MN106.** Genome-wide (left) and chromosome six (right) classification of sequence alignments between each genome and the MN106 reference genome from SyRI. Alignments are broken down into syntenic, not alignable to MN106, duplicated, inverted, inverted and translocated, and “highly-diverged region”.

##### Supplementary Figure 3 | Population differentiation by genomic class

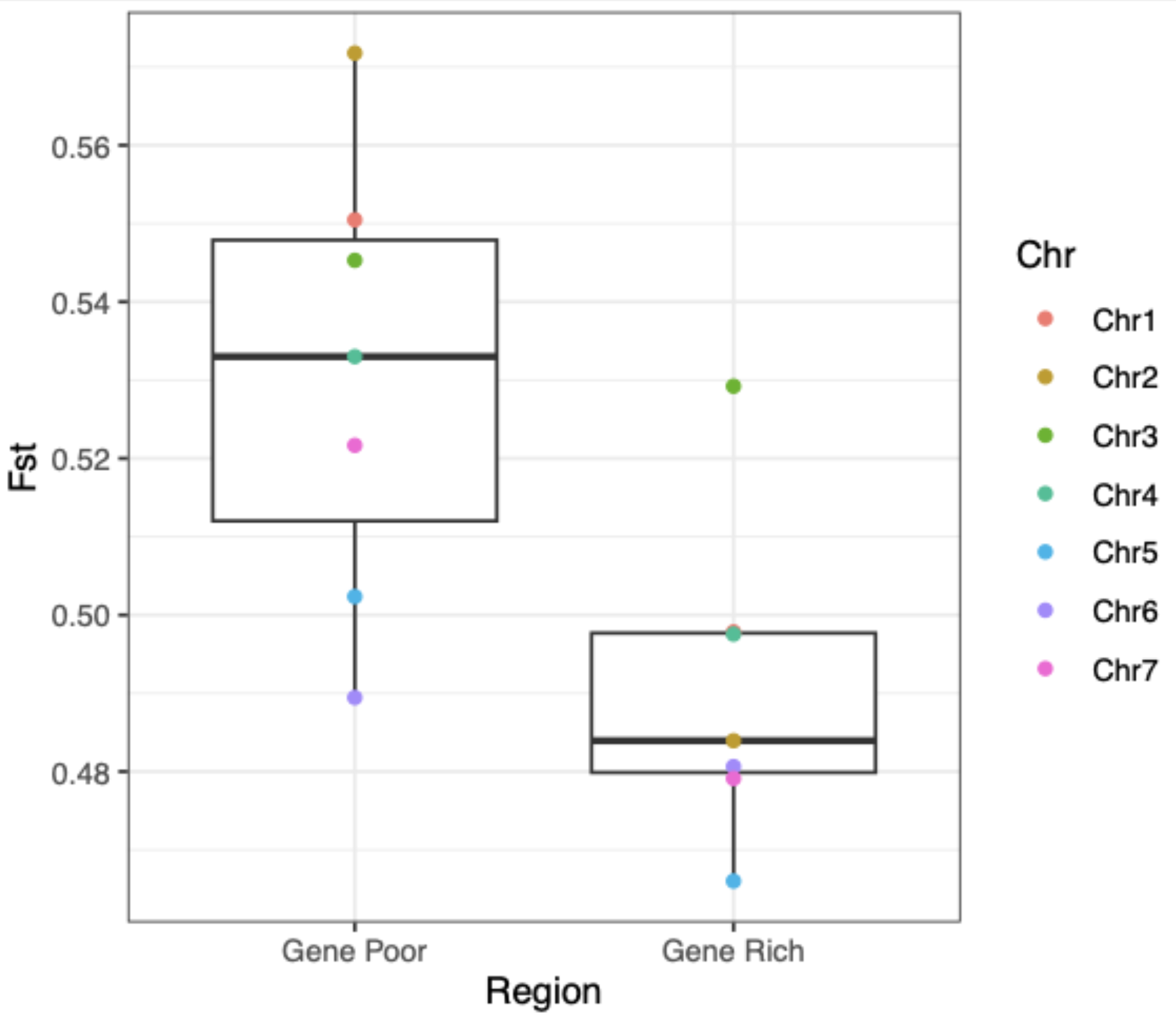

**Supplementary Figure 3 | Population differentiation by genomic class.** Comparison of Fst values (Hudson et al. 1992) between Armenian and non-Armenian accessions for each chromosome. Fst was calculated independently for SNPs located in gene-poor peri- and centromeric regions and gene-rich chromosome arms.

##### Supplementary Figure 4 | Position and structure of centromeric repeats

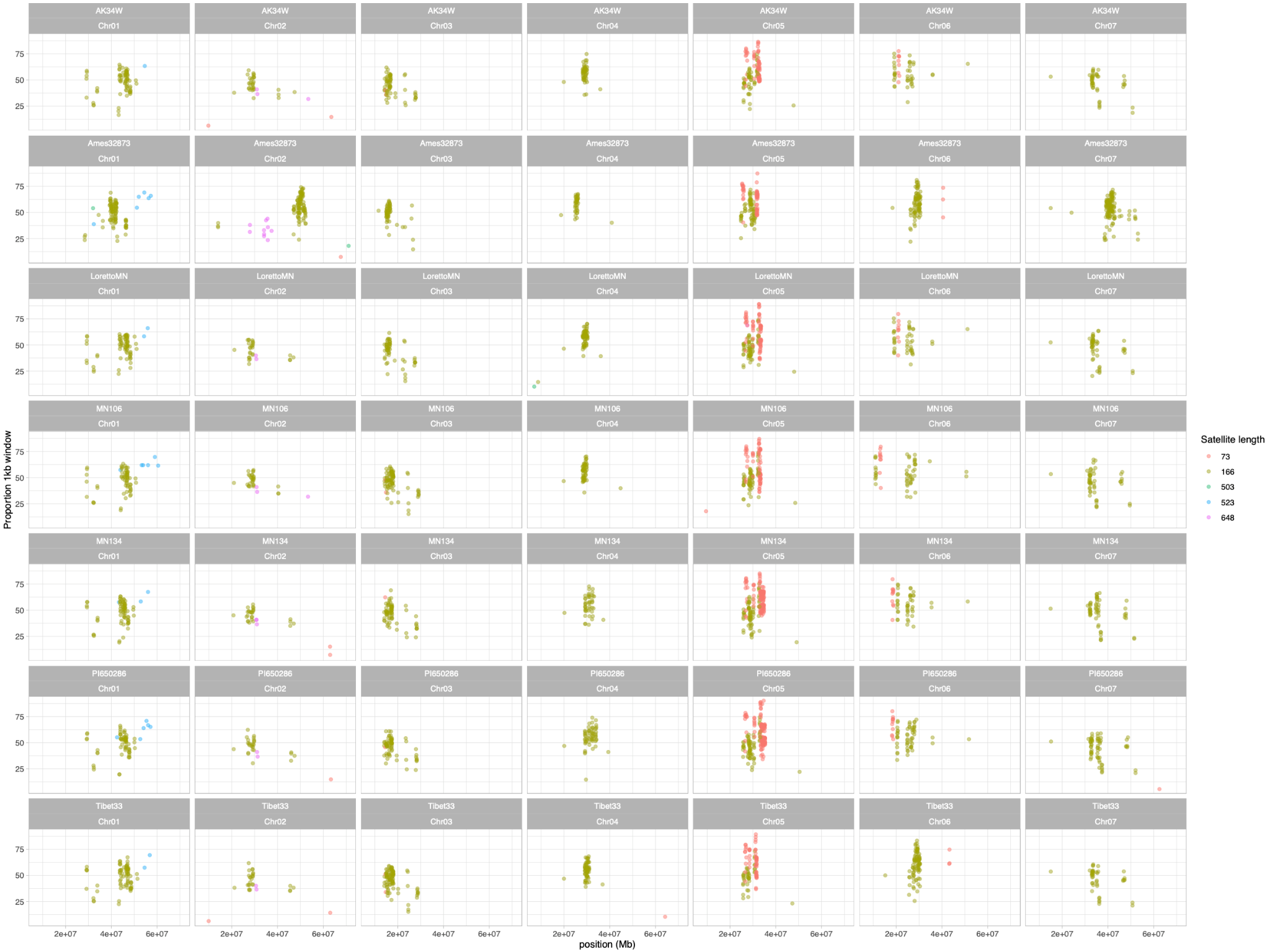

**Supplementary Figure 4 | Position and structure of centromeric repeats.** Content of all five repetitive sequences identified by TRASH across all seven genomes. Axes: genomic position, proportion of 1kb window consisting of putative satellite. The 166-bp and 73-bp satellites were incorporated into downstream analyses; the 523- and 503-bp sequences showed partial similarity to 16sRDNA.

#### Supplementary Tables

##### Supplementary Table 1 | NLR gene and cluster counts by accession.

**Supplementary Table 2 | NLR gene and cluster counts by accession.** Number of NLR genes identified, number of clusters, and cluster size for the seven pennycress genomes. Clusters were defined as at least two genes annotated as NLRs within no more than 50kb of each other.

| **Accession** | **Total Number of NLR genes identified** | **Number of NLR clusters** | **NLR cluster size range** |
| --- | --- | --- | --- |
| AK34W | 142 | 30 | 2-6 |
| Ames32873 | 143 | 30 | 2-8 |
| Loretto | 131 | 28 | 2-6 |
| MN106 | 141 | 28 | 2-7 |
| MN134 | 148 | 28 | 2-11 |
| PI650286 | 136 | 26 | 2-10 |
| Tibet | 145 | 33 | 2-6 |

##### Supplementary Table 2 | NLR cluster syntenic block overlap.

**Supplementary Table 2 | NLR cluster syntenic block overlap.** Although NLR clusters occupy only a small proportion of the genome, regions containing pennycress NLR clusters are extremely likely to be syntenic to regions containing Arabidopsis NLR clusters. Syntenic blocks between Arabidopsis and each pennycress genome were extracted from ‘synHits.txt’ in GENESPACE output. Overlap was calculated using custom R scripts.

| **Accession** | **Total number of syntenic blocks with Arabidopsis** | **Number of syntenic blocks containing an arabidopsis NLR cluster (percent of total blocks)** | **Base pairs containing by NLR clusters / total scaffold length (percent of total length)** | **NLR clusters in syntenic blocks containing arabidopsis NLR cluster / total NLR clusters (percent of total clusters)** |
| --- | --- | --- | --- | --- |
| AK34W | 134 | 21 (15.7%) | 1,099,594 / 456,034,515 (0.2%) | 30/30 (100%) |
| Ames32873 | 128 | 18 (14.1%) | 925,322 / 460,145,399 (0.2%) | 29/30 (96.7%) |
| Loretto | 133 | 18 (13.5%) | 923,847 / 455,918,564 (0.2%) | 28/28 (100%) |
| MN106 | 139 | 22 (15.8%) | 975,682 / 455,918,564 (0.2%) | 27/28 (96.4%) |
| MN134 | 140 | 20 (14.3%) | 969,396 / 461,649,255 (0.2%) | 27/28 (96.4%) |
| PI650286 | 138 | 20 (14.5%) | 756,104 / 466,533,064 (0.16%) | 24/26 (92.3%) |
| Tibet | 128 | 18 (14.1) | 975,908 / 456,735,805 (0.2%) | 31/33 (93.9%) |
