## Supplementary figures and images for "Structure and sequence evolution in the pennycress (*Thlaspi arvense*) pangenome"

### Supplementary Fig. 1

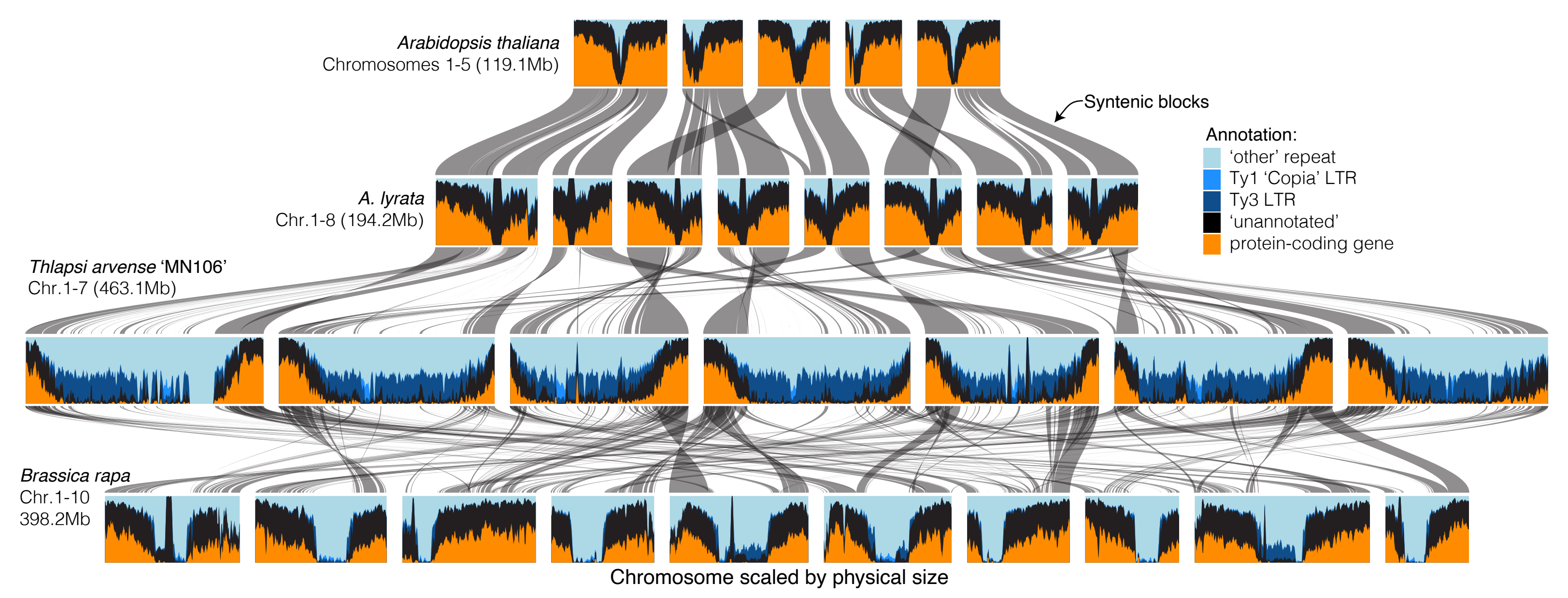

### Supplementary Fig. 2

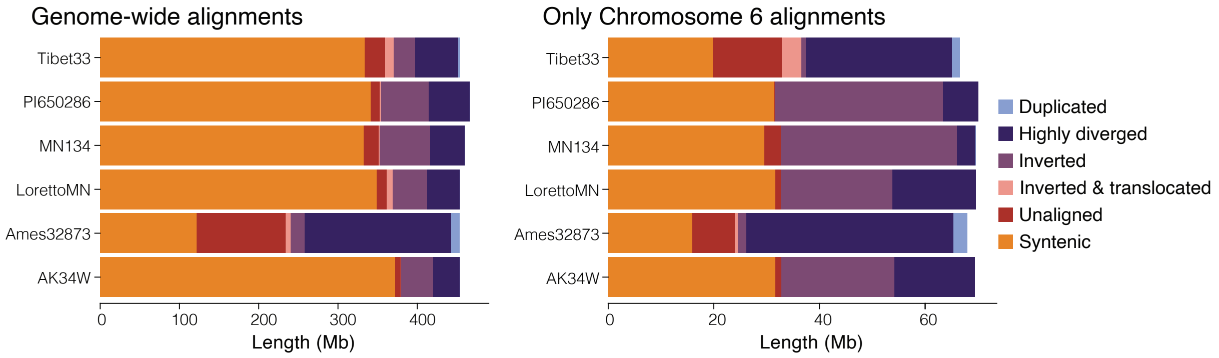

### Supplementary Fig. 3

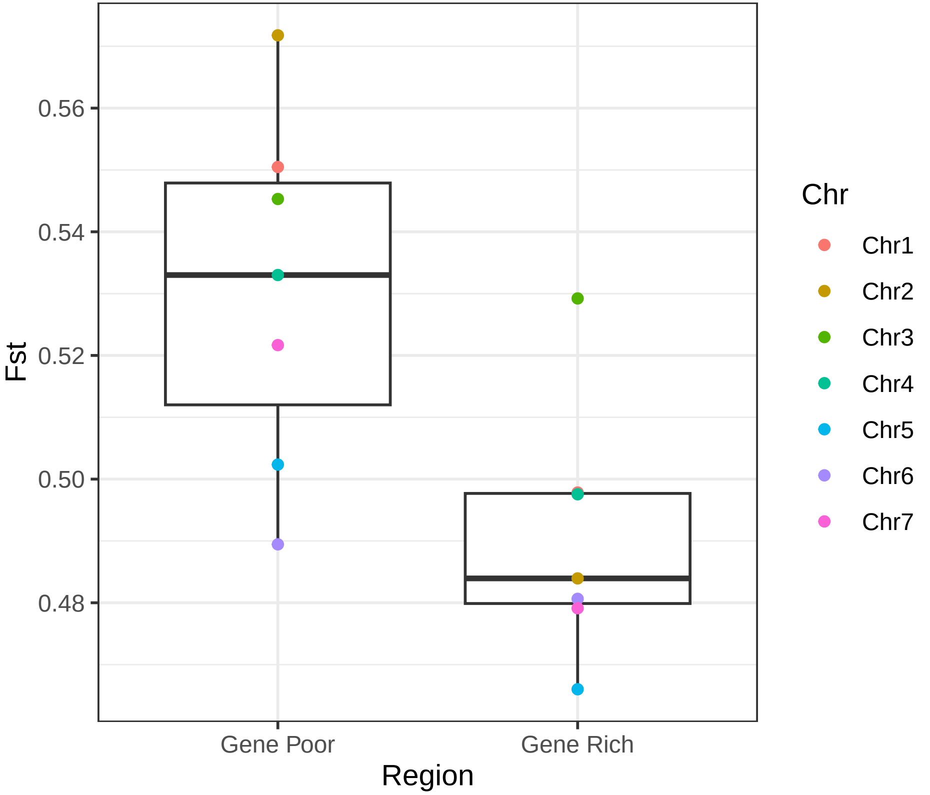

### Supplementary Fig. 4

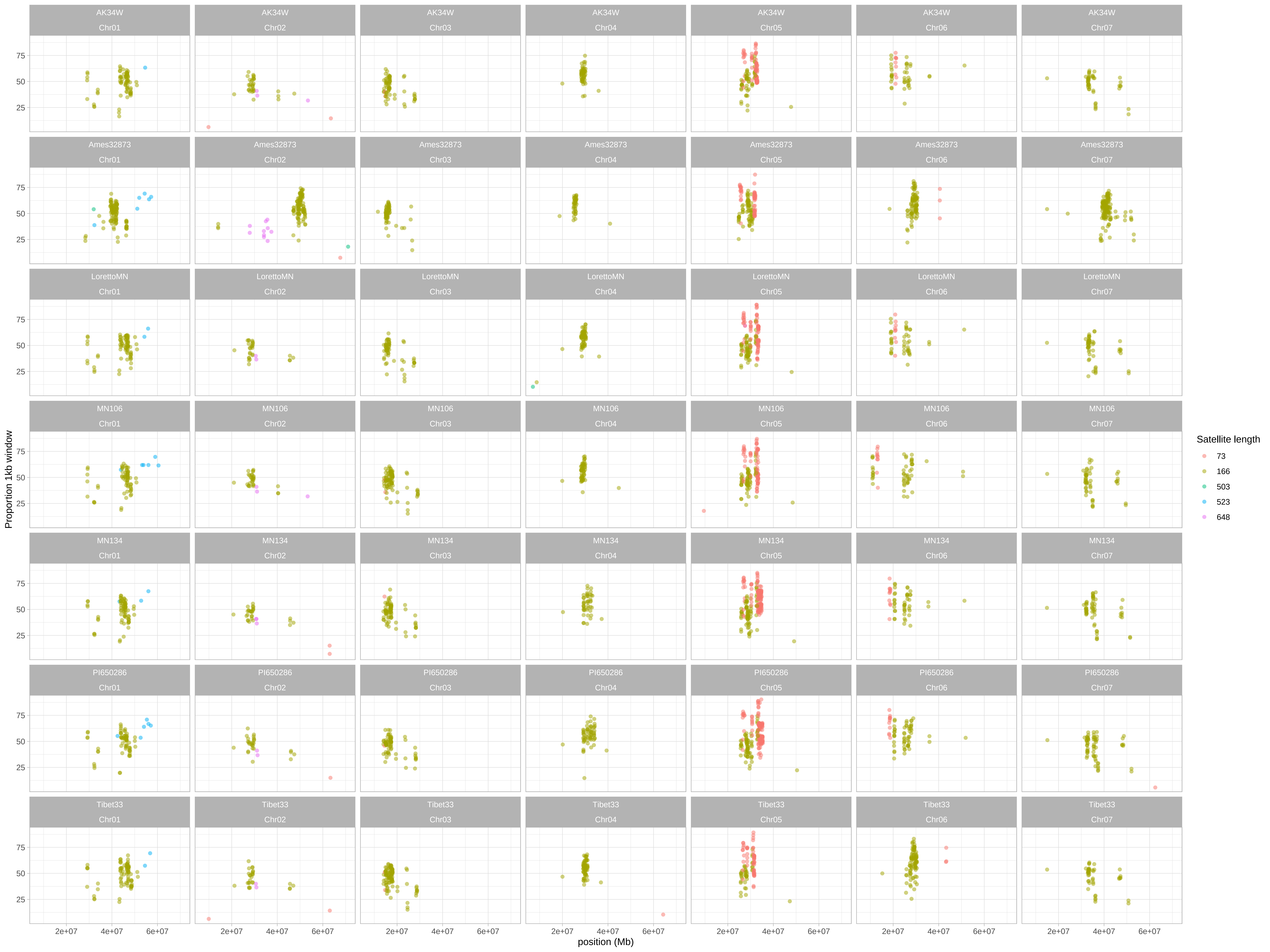
